# Normalized Raman Imaging for Studies of Tissue Physiology of the Kidney

**DOI:** 10.1101/2025.10.12.681746

**Authors:** William V. Trim, Seungeun Oh, Mariia Diakova, Kseniia Petrova, Takaharu Ichimura, Ayumi Takakura, Ranit Karmakar, Simon F. Nørrelykke, Leonid Peshkin, Joseph V. Bonventre, Marc Kirschner

## Abstract

Conventional histological relies on fixation, embedding, sectioning, and staining methods that distort cellular architecture, extract lipids, and introduces variability, limiting reproducibility. We present Normalized Stimulated Raman Imaging (NoRI) that enables quantitative, label-free protein and lipid measurements at high-spatial resolution. NoRI computationally corrects protein, lipid, and water Raman signals, allowing quantitative biomass measurements while preserving tissue architecture, facilitating analysis by convolutional neural networks. Applied to mouse kidney, NoRI accurately classified tubule types (F1-[harmonic mean of precision and recall against manual annotation]=0.93), anatomical regions (F1=0.91), and biological sex (F1=0.97), revealing greater cytoplasmic lipid (+6.9mg/mL; p=0.028) and nuclear protein (+26.3mg/mL; p<0.001) in female tubules. In acute kidney injury, NoRI captured dynamic lipid and protein organization and quantified brush border remodelling and lipid droplet changes over 25 days. Spatial lipid quantification was central for feature classification in AKI (F1=1.0). These results establish NoRI as a reproducible, quantitative method for diagnostics, histopathology, and feature discovery.

## Introduction

Following the invention of the compound microscope by the Janssens in the late 16^th^ Century, Marcello Malpighi and other physicians pioneered its use for the study of the micro-anatomy of organs(1). By the 19^th^ Century, the modern field of histology had been born. For the succeeding two centuries, microscopy has been at the center of discovery in the biological and medical sciences and for the routine diagnosis of many pathologies, especially cancer. During this time, there were significant improvements in microscopes themselves, but arguably histopathology depended on parallel developments of highly effective means of fixation, embedding, sectioning, and staining. Today’s standard approaches to modern histology are closely related to the technologies developed during the golden age of histology in the latter 19^th^ Century(2). Such tools and methods were responsible for some of the greatest discoveries in biology and embryology sciences in the 19^th^ and 20^th^ Centuries. Today, routine clinical pathology still makes use of many of these same approaches, such as formaldehyde as a fixative, paraffin embedding and mechanical sectioning to visualize tissues, and hematoxylin and eosin staining, devised by Nicolaus Wissotsky in 1877. For a long time, it has been appreciated that these means of tissue preparation, though revealing discrete highly contrasting details of cell structure, such as of chromosomes on the mitotic spindle, nevertheless distort more subtle features, which are blurred by the harsh chemical treatments, which also distort cellular structures and extract components of living cells, such as lipids and water-soluble materials. There is also considerable variability in the methods themselves due to variations in dye batches and inconsistent lab practices. Compensating for this inconsistency and distortion is difficult, since no gold standard for comparison exists. Nevertheless, histopathology remains an important part of the armamentarium of both clinical practice and fundamental biology of multicellular animals.

To address the limitations of conventional histological approaches, particularly the complete absence of lipid information, we have recently developed a Normalized (stimulated) Raman imaging microscope with associated methods (NoRI)(3, 4). Stimulated Raman scattering microscopy (SRS) had previously been used to quantify cellular materials by Freudiger*, et al.* (5), particularly to reveal lipids but its quantitative limitation by heterogeneous signal attenuation in thick tissue samples limited its applicability(6). To overcome this, Oh*, et al.* (3) developed a normalization process that relied on an additional channel for water. NoRI enabled precise determination of cell protein and lipid mass, at high spatial resolution, in live or fixed tissues by computationally removing light scattering artefacts. Furthermore, it was highly quantitative, a feature important for application of machine learning. With NoRI, chemical composition (namely, protein [CH_3_ bonds], lipid [CH_2_ bonds], and water) is automatically converted into absolute concentrations(3). This allows for absolute quantification of cellular protein, lipid, and water; such calculations enable extraordinarily accurate cross-comparisons of samples for feature identification. Consequently, NoRI data are unique not only due to their quantitative nature or ability to visualize lipids but also owing to their un-disturbed view of preserved structures. Nevertheless, detailed histopathology has not been explored in complex tissue specimens using NoRI to date.

For performing a histopathological analysis using NoRI, we selected the kidney as a candidate organ. The primary functions of the kidney, which Claude Bernard credited with maintaining the “milieu interieur,” include: 1. Blood filtration for waste removal, 2. Regulation of total body fluid balance, 3. Metabolic blood acid-base balance, and 4. Regulation of blood pressure. The kidney is also optimal for testing the suitability of NoRI for histopathology owing to its diverse array of distinct cell types comprising the functional unit, or nephron, which has many subunits, as well as the vasculature and interstitium. The kidney is also ideal since it is composed of distinct anatomical regions, including cortex, and outer and inner medulla. Given the critical connection of the kidneys’ own physiological function to whole-body physiology, understanding of the microanatomy of the kidney is vital to understanding both its physiology and how it regulates the physiology of the organism. More recently, machine learning methods have attempted to advance kidney histopathology and have offered the prospect of assisted diagnostics for clinicians and pathologists(7-9). Nevertheless, these methods are severely limited by the high variability in the primary image data derived using traditional techniques. As such, deep-level information related to lipids and microarchitecture is lost, and inconsistency is introduced between samples, making the generation of robust models and discovery of generalizable features difficult. Conventional histology, with its limited quantitative capacity and omission of lipids, may never be able to recover subtle, detailed information needed for deep machine learning models that would allow reliable diagnosis of kidney pathology.

As a first step to articulate a quantitative and minimally perturbed analysis of kidney microanatomy we utilized quantitative, highly sensitive image data obtained from NoRI microscopy in healthy and diseased mouse kidneys. The consistency of the new microscopic methods enabled an unprecedented application of machine learning, based on absolute quantification of the protein and lipid signals. Application of convolutional neural networks, a powerful form of machine learning, to the NoRI images in the setting of kidney ischemia-reperfusion injury—a model of acute kidney injury—allowed us to explore the potential sensitivity and reproducibility of NoRI for its application to histopathology and feature discovery. These findings show how NoRI represents a unique tool to explore unknown biological phenomena and provide a basis for advanced, automated histopathology in complex live or fixed tissues.

## Methods and Materials

### Animals

Experiments on kidney anatomy and sex differences were performed on C57BL/6 mice, purchased from Charles River Laboratories, which were housed at the Harvard Institutes of Medicine animal facility. Unilateral ischemia reperfusion experiments were performed in male BalbC mice (*described below*). Sixteen-week-old mice weighing 23.0–30.6 g (mean: 27.3 g) were used throughout. All animal protocols were approved by the institutional animal care and user committee of the Harvard Medical School.

### Unilateral ischemia reperfusion surgery

Studies were performed according to the animal experimental guidelines issued by the Animal Care and Use Committee at Brigham and Women’s Hospital, Boston, USA. Male BalbC mice were obtained from Charles River Laboratory and were allowed free access to water and standard mouse chow. Animals were anesthetized with pentobarbital sodium (60mg/kg body weight, intraperitoneally) prior to surgery. Analgia was provided with Ethiqa XR (3.25mg/kg). The mice were placed on a heating pad and body temperatures were maintained between 36.8-37.2°C throughout the procedure. The kidney was exposed via flank incisions, and unilateral renal ischemia was induced by clamping left renal pedicle with non-traumatic microaneurysm clamps (Roboz, Rockville, MD, USA) for 20 min. The kidneys were returned to the original place, and the incisions were temporarily closed during ischemia. After the clamps were removed, reperfusion of the kidney was visually confirmed. The muscle layer was closed using absorbable sutures and the skin incision was closed using sterile staples. One millilitre of pre-warmed 0.9% normal saline was administered after surgery. The mice were kept on a warm pad until fully recovered, then returned to their cages. Moist food pellets were placed on the bedding for easy access, and animals were monitored twice daily.

### Kidney resection and fixation

Following sacrifice, mice were perfused with chilled (4°C) phosphate buffered saline (PBS) containing 10 U/mL sodium heparin. Kidneys were then resected and cut in half, placed into 5 mL 4 % formaldehyde in PBS (Thermo-Fisher Scientific; MA, US), and stored at 4°C for 24 hours. Following fixation, samples were washed twice with chilled PBS and were stored in PBS containing 0.02 % sodium azide (Sigma-Aldrich; MilliporeSigma. MA, US) at 4°C for no longer than one-week before analysis.

### Kidney slices and immunofluorescence processing

Whole-kidneys were removed from storage and placed into mounting cylinders compatible with the Precisionary ZF-310-0F Compresstome® (Precisionary Instruments LLC.; MA, US) containing 2 % agarose diluted in distilled water (pre-heated to 50°C) (EZ Pack Agarose™; Benchmark Scientific, Inc.; NJ, US). All buffers used were filtered through a 0.22 µm filter (Corning®; AZ, US) prior to use. Constructs were then kept at 4°C for 30 minutes to harden the agarose-sample construct. Samples were next placed into the Compresstome® and cut into 120 μm slices at speed setting 4, oscillation setting 2. Individual tissue slices were then placed into ∼2 mL chilled PBS (0.02 % sodium azide) within a 12-well plate (VWR International, LLC; NJ, US). Samples were then placed into PBS containing 5 % normal donkey serum (Jackson ImmunoResearch Laboratories Inc.; PA, US) containing 0.02 % sodium azide overnight at 4°C with gentle rocking. Blocking solution was removed and replaced with PBS containing 0.02 % sodium azide at room temperature with gentle rocking for 20 minutes, repeated three times; these procedures were found to have no influence on the quantification of protein or lipid by NoRI (**Supplementary Fig. 1**).

After washing, the wash buffer was replaced with primary antibodies in tandem in PBS containing 0.02 % sodium azide for combinations of the following antibodies; Rat anti-mouse endomucin (1: 200) (sc-65495; Santa Cruz Biotechnology, Inc.; CA, US), sheep anti-mouse uromodulin (1: 200) (T0850-02B; USBiological Life Sciences; MA, US), and rabbit anti-mouse aquaporin 2 (1: 200) (29386-1-AP; Proteintech, ChromoTek GmBH; IL, US). Samples were incubated in primary antibody cocktail for 24 hours at room temperature with gentle rocking. Primary antibody cocktail solution was then removed and replaced with PBS containing 0.02 % sodium azide for 30 minutes at room temperature with gentle rocking to wash. This wash was repeated 4–6 times.

For samples where lotus tetragonolobus lectin (LTL) labelling of brush border was performed following primary antibody incubations, tissue sections were placed into PBS spiked with 10 μM calcium chloride and avidin blocking solution (Vector Laboratories; CA, US) according to manufacturer recommendations for 30 minutes at room temperature with gentle rocking. From this point on, all buffers were spiked with 10 μM calcium chloride according to manufacturer recommendations for samples where LTL was labelled. Samples were then washed at 20-minute intervals for 60 minutes total in PBS containing 0.02 % sodium azide. After washing, samples were placed into PBS (+10 μM calcium chloride) and biotin blocking solution (Vector Laboratories; CA, US) according to manufacturer recommendations for 30 minutes at room temperature with gentle rocking. Samples were then washed at 20-minute intervals for 60 minutes total in PBS containing 0.02 % sodium azide. Samples were then placed into biotinylated LTL antibody solution (1: 250) (B-1325-2; Vector Laboratories, Inc.; CA, US) in PBS (+10 μM calcium chloride) containing 0.02 % sodium azide for 24 hours at room temperature with gentle rocking. Following LTL incubation, samples were washed in PBS (+10 μM calcium chloride) containing 0.02 % sodium azide for 60 minutes 4–6 times at room temperature with gentle rocking. Next, samples were incubated in secondary antibody cocktail solution for 24 hours at room temperature with gentle rocking, in darkness, in PBS (+10 μM calcium chloride) containing 0.02 % sodium azide with combinations of the following antibodies; AffiniPure donkey anti-sheep Alexa Fluor® (AF)488 (1: 250) (Jackson ImmunoResearch Laboratories Inc.; PA, US), AffiniPure donkey anti-rabbit AF647 (1: 250) (Jackson ImmunoResearch Laboratories Inc.; PA, US), AffiniPure donkey anti-rat DyeLight™ (DL)405 (1: 250) (Jackson ImmunoResearch Laboratories Inc.; PA, US), and streptavidin AF594 (1: 250) (ThermoFisher Scientific, Inc.; MA, US). Secondary antibody cocktail solution was removed and replaced with PBS (+10 μM calcium chloride) containing 0.02 % sodium azide for 30 minutes at room temperature, in darkness, with gentle rocking, repeated 4–6 times.

After washing, samples were mounted onto glass slides (Superfrost® Plus VWR MicroSlides; VWR International LLC.; PA, US) with a small amount (∼75 μL) of anti-fade mounting solution containing 0.2 % propyl-gallate (Sigma-Aldrich; MA, US) diluted in 0.8 % fully deuterated dimethylsulfoxide (DMSO; MilliporeSigma; MA, US) in PBS (+10 μM calcium chloride) containing 0.02 % sodium azide. The deuterated form of DMSO was used so as not to contribute to the NoRI quantitation of the protein methyl groups. Number 1.5 micro cover glass slides (VWR International, LLC.; PA, US) were used to cover samples and slides were sealed using clear nail polish. Samples were stored at 4°C in darkness in a humidified container until imaging.

### Fluorescence microscopy and normalized Raman imaging (NoRI)

Microscopy imaging and quantitation was performed on an Olympus FV3000 using Fluoview (Olympus; Tokyo, Japan) and MatLab™ (MathWorks; CA, US). Immunofluorescence imaging was used for cell-type and tubule identification and for identification of markers of cellular stress. The immunocytochemistry was followed by NoRI imaging for protein and lipid quantification. All images were taken using a water immersion 60x 1.2NA objective lens (Evident-Scientific) and an oil immersion 1.4NA condenser lens, where each pixel was equivalent to 414nm for both NoRI and immunofluorescence imaging.

Protein and lipid concentrations were calculated from SRS images using NoRI as previously described(3, 4). Briefly, SRS images at 2853 cm^−1^, 2935 cm^−1^, and 3420 cm^−1^ bands (corresponding to the methylene- and methyl-groups, and water characteristic vibrational bands, respectively) were acquired from live or fixed cells using a custom-built spectral-focusing femtosecond SRS microscope. This microscope was constructed using synchronized femtosecond pulse lasers for the Pump and Stokes beams. A pair of dense flint (DF) glass rods chirped the pulses. An electro-optical modulator (EOM) modulated the amplitude of the Stokes beam at 20 MHz. A retro-reflector prism mounted on a motorized delay was adjusted to control the overlap of the Pump and Stokes beams. The Pump and Stokes beams, combined by a dichroic mirror (DM), were focused on the sample by the objective lens. Images were acquired by point-scanning using a pair of galvanized mirrors (scan mirror). After passage through the stimulated Raman scattering at the sample plane, the Pump beam was collected by a high numerical aperture condenser lens, selected by a short pass filter (SF), and its intensity was measured by a high-speed photodetector. The three SRS images were spectrally unmixed into protein, lipid, and water components using reference spectra measured from bovine serum albumin solution in water, dioleoyl-phosphocholine solution in per-deuterated methanol, water, and per-deuterated methanol. Unmixed images of protein, lipid, and water components were converted to the absolute concentration by using the sum of the three components as the normalization reference at each pixel. A representative region within each kidney that were imaged is presented in **Supplementary Fig. 2**.

### Data analysis and image segmentation

All images were processed by customized codes in Matlab™ (Mathworks) or ImageJ and FiJi (National Institute of Health). NoRI and immunofluorescence images were stitched together using the Grid/Collection stitching plugin for Fiji(10), and stacked into one multi-channel image using a custom macro for FiJi.

This study utilized an automated pipeline to quantify protein and lipid concentrations within the cytoplasm of individual kidney tubules. The approach involved three key steps: segmentation, classification, and measurement. A detailed description of this methodology, along with its validation and application, is also available in Karmakar, Trim, Kirschner and Noerrelykke (11). Segmentation was performed using a combination of classical image processing, deep learning and machine learning techniques, including the Segment Anything Model (SAM) for tubule segmentation(12); with the exception of glomeruli, which were hand-segmented. Post-processing steps, such as morphological operations and expert validation, were applied to refine segmentation accuracy. For comparison with SAM, a YOLOv8-based object segmentation model(13) was also fine-tuned to detect kidney tubules in NoRI images using annotated data derived from the protein and lipid signals. Fine-tuning was conducted on 640×640-pixel image crops, batch size of 16 with 200 epochs (iterations). Early stopping (where learning ends once a model’s metrics do not continue to improve further) after 30 epochs was performed. Data augmentation was applied to ensure robustness, using horizontal/vertical flips, random rotations, brightness/contrast adjustments, and affine transformations. All images were normalized using standard ImageNet mean and standard deviation values(14). The model was configured for single-class detection and fine-tuned using the Ultralytics YOLO implementation(13). Tubule classification was performed using Ilastik’s(15) object classifier, leveraging immunofluorescence markers (LTL, UMOD, and AQP2) to distinguish tubule types.

Nuclei were segmented using a U-Net model fine-tuned on manually labelled binary masks(16). The input data consisted of 640×640 patches from the protein channel, and corresponding nuclei masks were manually annotated based on visible nuclear morphology. The U-Net architecture followed a standard encoder–decoder design with skip connections and was implemented in PyTorch(17). Fine-tuning was conducted for up to 200 epochs, with a batch size of 8, and learning rate of 0.0001, where an epoch represents one full pass through the dataset, batch size represents the number of samples processed before the model updates its weights, and learning rate represents a step size for adjusting model weights during optimization. For fine-tuning, we used the binary cross-entropy loss and the Adam optimizer(18). Early stopping was applied after 30 epochs without validation improvement. The training data underwent geometric (i.e., rotations, cropping, scaling changes) and photometric (i.e., altered brightness/contrast, noise addition, blurring) augmentations, and all images were normalized to ImageNet statistics. We chose the model with the best Dice score—where Dice score represents the similarity between two segmented polygons, calculated as twice the intersection area divided by the sum of the areas of the two polygons—as a final model(16). Brush borders were segmented using a U-Net model fine-tuned(16) on manually labelled binary masks. The input data consisted of tubule images of 128×128 patches from the protein channel and lipid channels, and corresponding brush border masks were manually annotated based on visible brush border morphology. The U-Net architecture followed a standard encoder–decoder design with skip connections and was implemented in PyTorch(17). Fine-tuning was conducted using the same parameters as for nuclei. To identify luminal regions within segmented tubules, hierarchical clustering to low-intensity pixels in NoRI images was applied. Pixels were extracted using intensity values below a predefined threshold for each segmented tubule from the maximum projection of all imaging channels. These pixels were clustered using agglomerative hierarchical clustering with a Euclidean distance threshold of 5 pixels. Clusters containing fewer than 75 pixels were discarded to reduce noise. The remaining clusters generated binary masks representing lumen locations within the tissue(19). Nucleoli were segmented using a U-Net model fine-tuned on manually labelled binary masks(16). The input data consisted of nuclei images of 32x32 patches from the protein channel and lipid channels, and corresponding nucleoli masks were manually annotated based on visible morphology. The U-Net architecture followed a standard encoder–decoder design with skip connections and was implemented in PyTorch(17). Fine-tuning was conducted using the same parameters as for nuclei.

Protein and lipid concentrations were quantified by measuring pixel intensities within the cytoplasm, obtained by subtracting the segmented nuclei, nucleoli, lumen, and brush border regions from the tubule mask. Intensity values were converted to concentrations using established conversion factors for protein and lipid. The resulting measurements provided insights into the spatial distribution of these biomolecules across different tubule types.

### Feature-based classification, feature extraction, and interpretability analysis

ResNet50 convolutional neural network(20), trained on ImageNet, was used to identify regions of interest in NoRI images. The model’s final fully connected layer (where every neuron is connected to every neuron in the neural network’s previous layer) was replaced with a new layer corresponding to the number of classes (e.g., cell types, sex, ischemic timepoint) in the dataset. Input images were organized into folders labelled by class and divided into training and validation sets. Image tiles were resized and randomly augmented through horizontal and vertical flips, rotations (0–270°), and random crops. All images underwent normalization using ImageNet’s mean and standard deviation. For model fine-tuning, we utilized cross-entropy loss and the Adam optimizer (learning rate = 0.0005) for up to 25 epochs, evaluating performance using accuracy and macro-averaged F1-score (a performance metric on an ascending scale of 0–1 for classification performance calculated as the harmonic mean of precision and recall [2/(precision^-1^ + recall^-1^]). The best-performing model weights (i.e., those closest to 1) were saved.

To extract high-level image features (determined by the neural network) and visualize class-specific patterns (i.e., features that determine the class) in NoRI images, fine-tuned ResNet50 models were employed and activations (the output signals produced by a layer of a neural network after it applies its transformation to the input) were retrieved from the penultimate average down sampling layer. Images were transformed using standard normalization and resizing procedures before being passed through the network in evaluation mode. The resulting feature vector was saved and used for dimensionality reduction and visualization with UMAP for each tile. In parallel, Grad-CAM (Gradient-weighted Class Activation Mapping(21)) was applied to layers 2, 3, and 4 of the ResNet architecture to produce class-discriminative heatmaps that highlight image regions most relevant to the model’s predictions. These heatmaps were overlaid on grayscale representations of the original inputs to facilitate biological interpretation. UMAP plots and Grad-CAM overlays were generated for validation samples to promote qualitative and quantitative assessment of class differences.

Following feature identification by ResNet50, to identify and quantify luminal regions and lipid droplets within segmented tubules and microvasculature regions, hierarchical clustering to low-intensity pixels in NoRI images was applied. Pixels were extracted using intensity values below a predefined threshold for each segmented tubule from the maximum projection of all imaging channels.

All code and materials used for image processing can be accessed at: https://github.com/HMS-IAC/nori, https://github.com/MDyakova/NORI_feature_extraction and https://github.com/MDyakova/NORI_image_segmentation. A schematic summary of segmentation and machine learning analysis pipelines is presented in **Supplementary Fig. 3**.

### Histological preparation and analysis

A single section of a kidney from five healthy male and five healthy female mice were fixed as described above and were dehydrated, paraffin-embedded, and stained with haematoxylin and eosin by INVIVOVEX LLC. (MA, US). Slides were imaged at 60x magnification on an Olympus slide view VS200.

### Statistical analysis

All statistical analyses and graphical representations were performed and produced using MatLab™ and GraphPad Prism (MA, US). MatLab scripts are accessible from: https://github.com/kirschnerlab/NoRI. Comparisons between variables, where normally distributed were performed using student’s *t-*tests. Where non-normally distributed, Mann-Whitney *U* non-parametric equivalents were used. Cumulative distribution analysis was performed using Kolmogorov-Smirnov tests. For analysis of time-course results, Analysis of Variance (ANOVA) tests were performed, with Bonferroni post-hoc corrections applied. For analysis of ≥3 groups, one-way ANOVA was performed, with Dunn’s post-hoc corrections applied. Distribution of raw protein and lipid concentrations between groups were compared using Kruskal-Wallis tests, with Dunn’s post-hoc corrections applied. Effect sizes were estimated using Cohen’s *d* calculations throughout. Specific uses of each statistical test are detailed in the respective figure legends. Statistical significance was set at *p*≤0.05.

## Results

To establish the potential of NoRI microscopy for exploring tissue biology we first needed to establish its accuracy in measuring proteins and lipids in a well-studied organ under varying conditions. After examining several organs, we chose the intact mouse kidney. We studied the kidney under three physiological or pathophysiological differences: female *versus* male, anatomical variation in the various tissues and cell types in the kidney itself, and the injury and recovery after ischemia-reperfusion injury induced by cessation of renal blood flow followed by reperfusion. At the same time, as we began to appreciate the power of this data set, we experimented with various machine learning methods for analysing NoRI images to measure changes in cells and tissues with known levels of confidence.

Samples were imaged by placement between a water immersion 60x 1.2NA objective lens (Evident-Scientific) and an oil immersion 1.4NA condenser lens, where each pixel equated to 414nm for both NoRI and immunofluorescence imaging. A representative kidney section imaged by NoRI is presented in **Fig. 1A**. A single tile (212µm^2^) image of NoRI and the corresponding immunofluorescence image is presented in **Fig. 1B**. From such images we derived a mask (i.e., segmented region of interest) of kidney tubules and other structures. A representative tubule structure and a corresponding sub-structure mask is also presented for this same region (**Fig. 1B**). Representative examples of the segmentation of nucleoli are presented in **Supplementary Fig. 4**, and representative images of all image channels and segmentation layers are presented in **Supplementary Fig. 5**. We found that we could fine-tune image models that would parse out tubules, lumen, nuclei, and brush border from the NoRI images, by using a combination of Segment Anything Module (SAM)(12) (for tubules) and YOLO(13) (all remaining sub-structures) fine-tuned against manual segmentation of tubules. Both SAM and YOLO had high mean average precision scores (a single metric that summarizes a model’s ability to correctly classify and localize specific objects [e.g., a tubule] across all classes and confidence thresholds, against a gold-standard [e.g., hand-segmented tubules]) for tubule and nuclei segmentation (**Fig. 1C**). We ascertained the variability in cytoplasmic protein and lipid concentrations in proximal tubules (LTL+), distal tubules (UMOD+), and connecting tubule and collecting ducts (AQP2+) in kidney slices from the Midpole region of eight kidneys from four male and four female C57BL/6 mice (**Fig. 1D**). We found that tubule types generally clustered based on their average cytoplasmic protein and lipid concentrations for collecting ducts (Protein: 138.5±19.7 and Lipid: 40.2±13.1mg/mL), proximal tubules (Protein: 194.3±18.4 and Lipid: 52.2±9.5mg/mL), and distal tubules (Protein: 170.8±16.1 and Lipid: 62.4±10.3mg/mL) (**Fig. 1D**). These results also demonstrated the similarity of tubule cytoplasmic concentrations across animals. Quantification of nuclear protein concentrations for different tubule types and brush border protein/lipid in proximal tubules, and protein and lipid concentrations within the glomeruli is presented in **Fig. 1E**. We observed that nuclei constitute 12.9±4.7% of the tubule area but contained negligible quantities of the lipid within the cell. We also found differences in nuclear protein concentration between tubule types that was statistically significant (134.7±19.3 *versus* 166.8±20.3 *versus* 130.7±22.4mg/mL for collecting ducts, proximal tubules, and distal tubules, respectively; *p*≤0.0001; **Fig. 1E**). When we took all segmented tubules (not segregated by sub-types), without immunofluorescence channels and processed them through ResNet50, the deep-convolutional neural network (CNN) pipeline, we found NoRI+CNN analyses had the strongest performance for tubule type prediction (mean ± SD F1-score: 0.93 ± 0.04; **Fig. 1G**). By comparison, NoRI protein/lipid data, geometric features (*listed in* **Supplementary Table 1**), or a combination of both were less powerful predictors of tubule type (F1-scores: 0.62±0.10, 0.58±0.13, and 0.67±0.12, respectively; **Fig. 1G**). The impressive UMAP representation of tubule types predicted by NoRI+CNN is presented in **Fig. 1H**.

**Figure 1.**
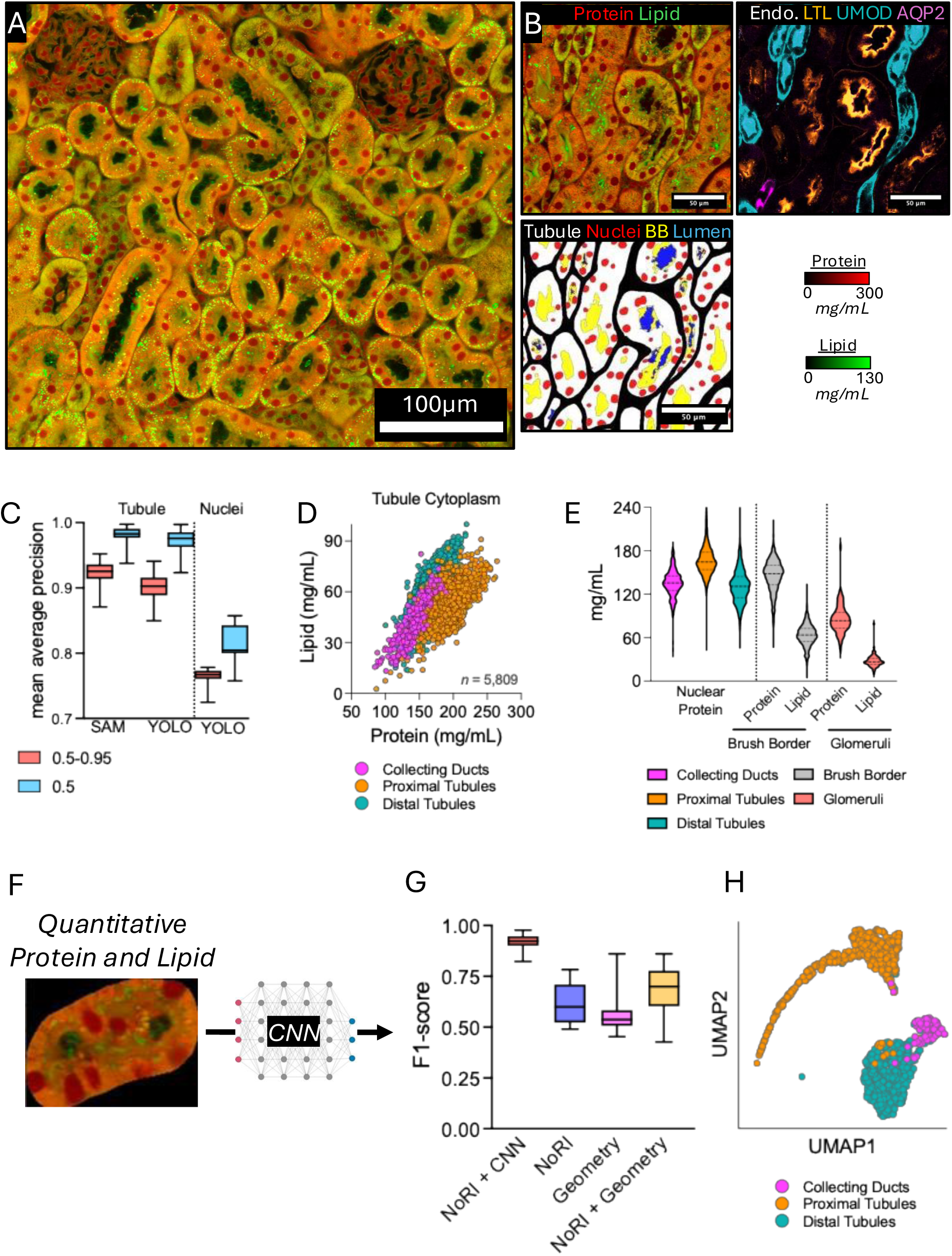
Overview of the use of NoRI and CNNs in mouse kidneys. A, Representative NoRI image of mouse kidney. Quantitative readouts of Protein (Red) and Lipid (Green) are displayed. Scale bars representing quantitative protein and lipid signals are presented to the right. B, Sub-section of mouse kidney simultaneously imaged by NoRI (Left) and immunofluorescence (Right) for tubule type identification, with a representative segmentation mask (Bottom) for tubules, nuclei, brush border, and lumen. C, Box-whisker plots of mean average precision (0.5 and 0.5-0.95) scores for tubules and nuclei using SAM and YOLOv.8. Scores are relative to hand segmentation of the same structures. Boxes represent median with 25^th^/75^th^ percentiles. Whiskers represent min-to-max values. D, Scatter plot of cytoplasmic protein and lipid content on a per-tubule basis for two (1x5 mm) kidney sections representing cortex to medulla from the Midpole portion of the left kidney. Color labels for tubule types were assigned according to immunofluorescence staining. E, Violin plot of nuclear protein, brush border, and glomeruli protein and lipid content on a per-tubule basis for tubules also displayed in Panel D. Data represent a total of 1,662 tubules. Violins are presented with median (dashed line) and 25^th^/75^th^ percentiles (dotted lines). F, Representative NoRI image of a single tubule analysed by ResNet50 for tubule-type identification by NoRI images without immunofluorescence. G, Box-whisker plots of F1-scores for tubules input into ResNet50 for tubule prediction. Comparisons between ResNet50 (CNN + NoRI) analyses is presented against each module within the data, including protein and lipid data alone (NoRI), Geometry only, and NoRI+Geometry. Boxes represent median with 25^th^/75^th^ percentiles. Whiskers represent min-to-max values. H, UMAP plot of tubules clustered by ResNet50. Color coding represents tubule identification assigned by immunofluorescence. Each datapoint represents an individual tubule.

Having found that different tubule types possessed different protein and lipid densities, we asked whether protein and lipid concentration of cells in the kidney vary depending on their anatomical location within the organ. To address this question, we took a section of kidney from three different anatomical regions, namely Rostral pole, Midpole, and Caudal pole portions of both left and right kidneys from two male mice and fine-tuned a ResNet50 pipeline on a sub-section of these images (**Fig. 2A**). The Caudal pole region had the highest F1-score for prediction (0.91±0.06; **Fig. 2B**), followed by the Rostral pole (F1-score: 0.79±0.14) and Midpole (F1-score: 0.58: 0.58±0.25) (**Fig. 2B**). We also identified statistically significant differences between all three anatomical regions for cytoplasmic protein, and between all regions, except for Rostral pole and Midpole (*p*=0.123; **Fig. 2E**), for lipid concentrations across tubules with highest cytoplasmic protein concentrations present in Caudal pole sections (Protein: 158.4±29.5 and Lipid: 49.9±11.5mg/mL), whereas Midpole and Rostral pole were similar across protein and lipid cytoplasmic concentrations ([Rostral pole] Protein: 134.6±21.0 and Lipid: 43.1±9.0mg/mL, and [Midpole] Protein: 136.6±20.8 and Lipid: 42.6±9.0mg/mL)(*ANOVA*: all *p*≤0.0001; **Fig. 2D**, **Fig. 2E** and **Supplementary Fig. 6**). These differences were further supported by medium-to-high effect sizes when comparing distinct tubule types between Rostral pole *versus* Caudal pole (*d*=0.63–1.21) and Midpole *versus* Caudal pole (*d*=0.88–1.76), but with less distinction between Rostral pole and Midpole portions (*d*=0.27–0.99) (**Supplementary Fig. 6**). We also observed a significant downward trend in proximal tubule cytoplasmic protein and lipid concentrations from cortex to medulla in the Midpole region (*r^2^*>0.281, *p*<0.0001; **Supplementary Fig. 7**). We did not identify any significant differences between regions from left and right kidneys (F1=0.58; *data not shown*).

**Figure 2.**
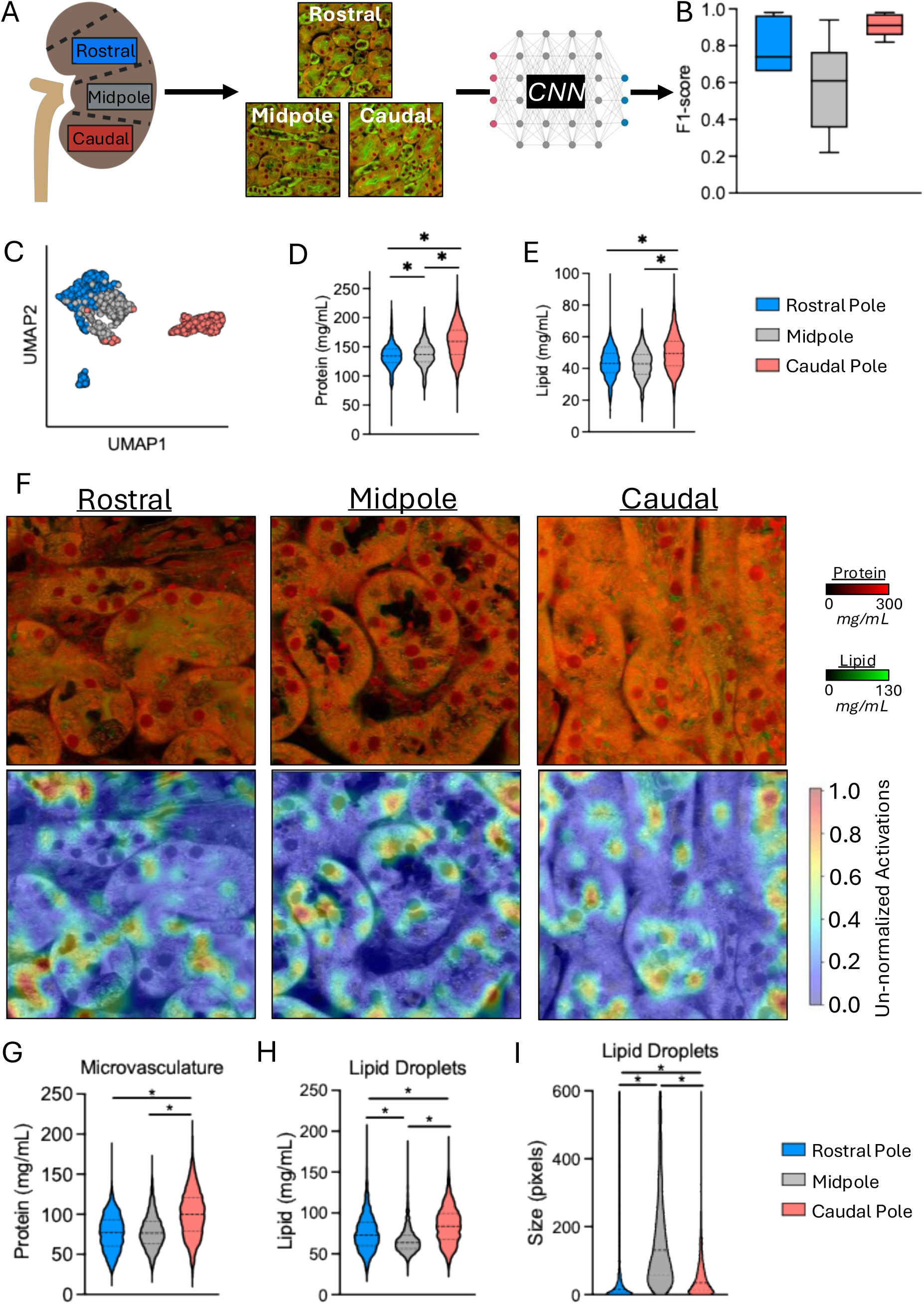
Exploration of kidney anatomy by NoRI. A, Schematic of experimental design, wherein 2-3 kidney sections from three distinct anatomical regions were imaged by NoRI and input into ResNet50. B, Box-whisker plots of F1-scores for anatomical region prediction on a per-tubule basis. Boxes represent median with 25^th^/75^th^ percentiles. Whiskers represent min-to-max values. C, UMAP plot of tubules color coded by anatomical region predicted by ResNet50. Each datapoint represents an individual tubule. D, Violin plots of tubule cytoplasmic protein content measured by NoRI, split by anatomical region. Violins are presented with median (dashed line) and 25^th^/75^th^ percentiles (dotted lines). E, Violin plots of tubule cytoplasmic lipid content measured by NoRI, split by anatomical region. Violins are presented with median (dashed line) and 25^th^/75^th^ percentiles (dotted lines). F, Representative NoRI images from each anatomical region with heatmap of features defining each region identified by ResNet50. G, Violin plot of microvasculature protein concentration identified by ResNet50 between anatomical regions. Violins are presented with median (dashed line) and 25^th^/75^th^ percentiles (dotted lines). H, Violin plot of lipid droplet lipid concentration identified by ResNet50 between anatomical regions. Violins are presented with median (dashed line) and 25^th^/75^th^ percentiles (dotted lines). I, Violin plot of lipid droplet size (in pixels) identified by ResNet50 between anatomical regions. Violins are presented with median (dashed line) and 25^th^/75^th^ percentiles (dotted lines). *Statistical comparisons in Panels D, E, G–I were performed using Kruskal-Wallis tests with Dunn’s post-hoc corrections applied*.

We used the ResNet50 pipeline fine-tuned on different kidney regions to identify structural features that might distinguish between regions (representative examples presented in **Fig. 2F**). These analyses identified protein concentrations in spaces between tubules where the tissue microvasculature resides—which we will term, “microvasculature.” These represent the microvasculature of the kidney and are a key distinguishing feature between regions, where the highest protein concentration was found in Caudal pole sections (100.5±29.4mg/mL; *p*<0.0001), while Midpole and Rostral pole were similar (77.2±22.3mg/mL and 78.3±21.2mg/mL, respectively; *p*>0.999)(**Fig. 2G**). We also found lipid droplet concentration and size were significantly different between regions (*p*≤0.0001). Specifically, Caudal pole exhibited highest lipid droplet concentration (84.7±22.9mg/mL), followed by Rostral pole (75.4±22.0mg/mL) and Midpole (66.5±15.6mg/mL)(**Fig. 2H**). Conversely, lipid droplet size was substantially larger in the Midpole (278±418 pixels) compared to Rostral or Caudal pole regions, which were similar, on average (61±193 and 67±98 pixels, respectively) (**Fig. 2I**).

After confirming that NoRI combined with CNNs can predict tubule type and anatomical location of tubules in a single kidney, we next turned our focus on examining sexual dimorphism, using four male and four female C57BL/6 mice. We analyzed the cytoplasmic protein and lipid concentrations to characterize broadly the differences between sexes (**Fig. 3A**). Cytoplasmic protein in LTL+ proximal tubules—specialized in resorption of substances back into the bloodstream—was significantly higher in female mice (256.9±6.9mg/mL), on average than for males (245.5±3.2)(*p*=0.029; **Fig. 3B**). Furthermore, cytoplasmic lipid content was between 6.5–7.3mg/mL more concentrated in female mice across all tubule types analyzed (*p*≤0.028; **Fig. 3C**). There were no differences between number of tubules between sexes (*p*≥0.767; **Supplementary Fig. 8**). To probe more deeply into the localization and features underpinning sex differences in mouse kidneys in NoRI images, we fine-tuned image tiles of varying sizes from 40–320 pixels in size (∼16.5–132.5µm^2^, ranging from a single cell to several tubules per image) to determine what type of input data in the ResNet50 pipeline best captured the differences (**Fig. 3D**). We found a tile size of 320 pixels (132.5µm^2^) achieved remarkable F1-scores for predicting the sex of a given tubule, likely due to capturing larger regions of tubules and other structures, though all image tile sizes worked to some degree, with high accuracy to predict the sex of a tubule (average F1-scores: 0.82–0.97; **Fig. 3E**, **Supplementary Fig. 9**). We next explored what were the actual features that ResNet50 that accounted for the sexual dimorphism (**Fig. 3G**). These analyses revealed that the level of nuclear protein and the relative abundance of protein to lipid in the microvasculature region were both significantly greater in females (+26.3mg/mL and +19.5%, respectively; *p*≤0.001; *d*=0.82–0.83; **Fig. 3H and I**). The concentration of lipid and the size of the lipid droplets were similarly significantly higher in females (+11.65mg/mL and +7.12µm^3^, respectively; *p*≤0.001; *d*=0.62–0.63; **Fig. 3J and K**). We also compared our ability to determine sex in the kidney with conventional histological methods (H&E staining) against our combination of NoRI plus machine learning analyses (**Supplementary Fig. 10**). To facilitate a direct comparison between H&E and NoRI images, we matched the input image data entirely. We used adjacent kidney sections (taken within 120µm of one another in the original kidney specimen) from the same four male and four female kidneys and used a total of 3,088 image tiles of 320x320 pixels at a resolution of 0.414µm/ pixel. As well as the same specimens and input data being used, we also used the same ResNet50 models for analysis of both H&E and NoRI images to directly compare their ability to distinguish male from female kidneys. We found that although sex could be reliably determined from H&E images using ResNet50 (presumably due to the relative importance of protein over lipid as shown by NoRI), the staining inconsistencies, lack of quantitative data, and disruption of tissue architecture in H&E staining failed to register finite/smaller (e.g., sub-cellular-sized) features that could distinguish males from females (i.e., at layers 2 and 3 of the ResNet50 output). Instead, activation maps presented in **Supplementary Fig. 10** highlight sex discrimination was largely focused on macro-structures such as entire tubule bodies, rather than sub-cellular structures/regions. This admittedly unusual metric suggests that NoRI can better distinguish subtle differences in structure than H&E staining and is much more quantitative, allowing for a direct measurement of such structures that are normally lost due to staining variability sample-to-sample, or micro-architectural disturbance from the extensive processing required by conventional histology protocols.

**Figure 3.**
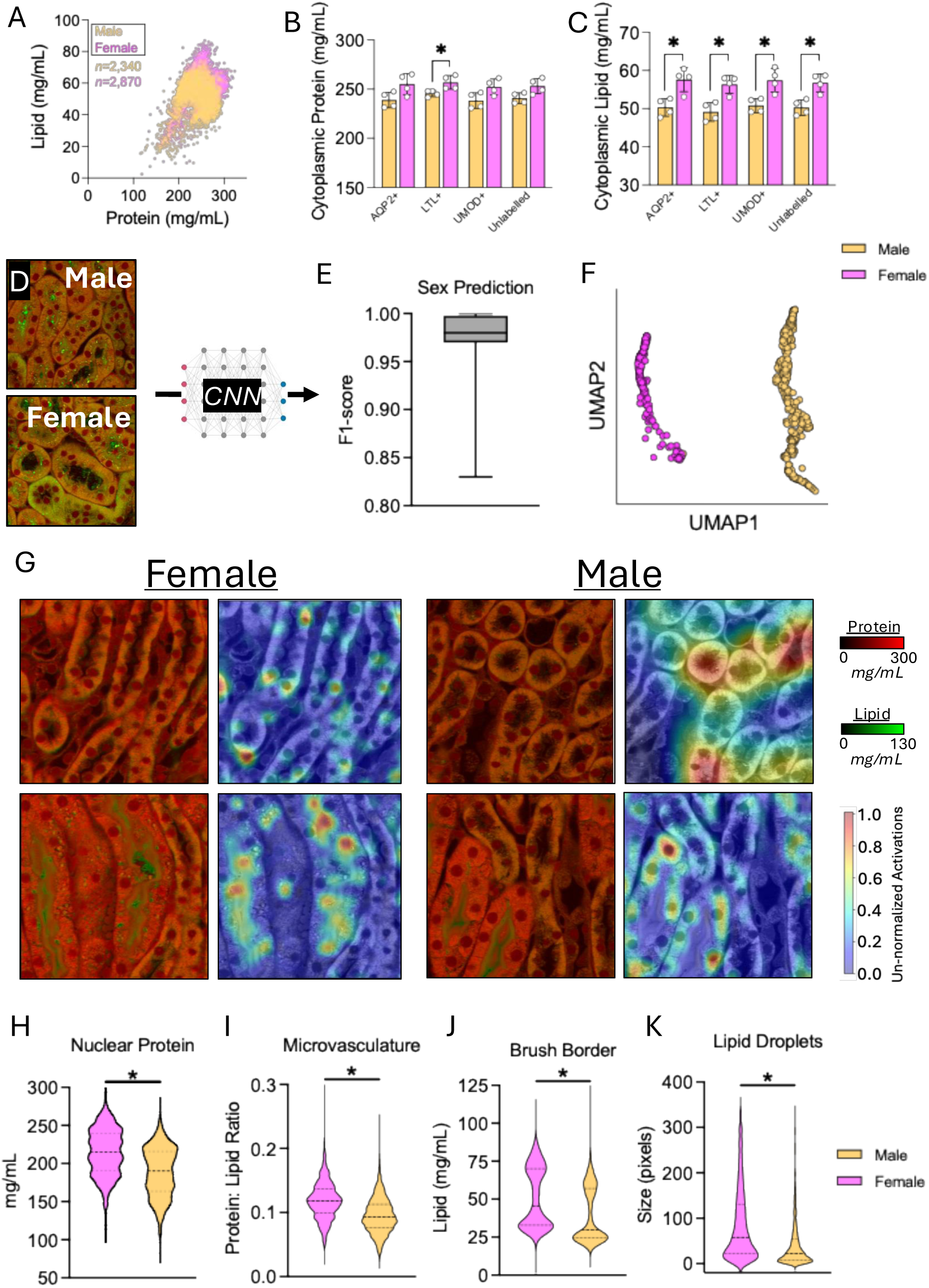
Exploration of kidney sex differences by NoRI. A, Scatter plot of tubule cytoplasmic protein and lipid content measured by NoRI in male and female kidneys (*n* = 4 mice per group). Each datapoint represents an individual tubule. B, Bar graph of cytoplasmic protein content, split by tubule type, measured by NoRI in male and female kidneys. Each datapoint represents the mean value across all tubules for an individual mouse. C, Bar graph of cytoplasmic lipid content, split by tubule type, measured by NoRI in male and female kidneys. Each datapoint represents the mean value across all tubules for an individual mouse. D, Representative NoRI images from male and female mouse kidneys input into ResNet50 for sex prediction on a per-tubule basis. E, Box-whisker plots of F1-scores for sex prediction on a per-tubule basis from NoRI images. Boxes represent median with 25^th^/75^th^ percentiles. Whiskers represent min-to-max values. F, UMAP plot of tubules color coded by sex and tubule type predicted by ResNet50. Each datapoint represents an individual tubule. G, Heatmap figures of female and male kidneys imaged by NoRI representing regions discriminating male from female kidneys according to ResNet50. H, Violin plot of nuclear protein concentrations identified by ResNet50 between sexes. Violins are presented with median (dashed line) and 25^th^/75^th^ percentiles (dotted lines). I, Violin plot of microvasculature protein: lipid ratio identified by ResNet50 between sexes. Violins are presented with median (dashed line) and 25^th^/75^th^ percentiles (dotted lines). J, Violin plot of brush border lipid concentration identified by ResNet50 between sexes. Violins are presented with median (dashed line) and 25^th^/75^th^ percentiles (dotted lines). K, Violin plot of lipid droplet size (in pixels) identified by ResNet50 between sexes. Violins are presented with median (dashed line) and 25^th^/75^th^ percentiles (dotted lines). *Statistical comparisons in Panels B and C were performed using Kruskal-Wallis tests with Dunn’s post-hoc corrections applied. Data in Panels H–K were analysed by Kolmogorov-Smirnov tests*.

Given the success of NoRI in predicting tubule type, anatomical location, and the sex of a tubule simply from structural and chemical composition (protein and lipid mass density) we applied NoRI and convolutional neural networks to better understand kidney dysfunction, in a model of transient unilateral ischemic-reperfusion injury (IRI). This model of acute kidney ischemia, caused by temporary occlusion of the renal artery and vein of one kidney, leaving the other as a contralateral control, is a common model for acute kidney injury (AKI)(22). When due to ischemia, AKI is characterized by oxygen deprivation to inflammation, oxidative stress, renal damage, and recovery which can be “adaptive” with few if any long-term consequences, or “maladaptive” resulting in incomplete repair with tissue scarring/fibrosis (22). As such, this model represents a powerful means of exploring both kidney damage and repair, as commonly occurs in human AKI.

In this AKI model, where occlusion occurred for only 20 minutes, we first performed a time course analysis of signs of dysfunction and repair across collecting ducts, distal and proximal tubules. We observed no change in tubule numbers between contralateral control and IRI kidneys at each time point (**Supplementary Fig. 11**). We next analysed the cytoplasmic protein and lipid concentrations of tubule types at each time point post-IRI and found that proximal and distal tubule cytoplasmic lipid concentrations showed changes during the recovery period post-IRI (+6.3mg/ml from day 1 to +8.3mg/mL at day 25 post-IRI, respectively). There were significant reductions in proximal tubule and distal tubule lipid concentrations between groups specifically at day 2 post-IRI (52.9±0.6mg/mL *versus* 46.2±0.2mg/mL, and 61.3±2.2mg/mL *versus* 47.0±3.3mg/mL for proximal and distal tubule cytoplasmic lipid concentration in contralateral control *versus* IRI kidneys at day 2 post-IRI, respectively; two-way ANOVA: *p*≤0.037; **Supplementary Fig. 12**). We next explored the time course relative to the contralateral control. We took NoRI images from days 1, 2, 5, and 25 post-IRI for both contralateral control and ischemic kidneys and followed the time course of structural changes in kidney dysfunction and repair following the ischemic injury (**Fig. 4A**). In support of the above findings in **Supplementary Fig. 12**, we found the greatest difference between contralateral control and IRI (F1-score: 0.97±0.04) at day 2 post-IRI (**Fig. 4B**). Days 1 and 5 post-IRI displayed similar degrees of separation (F1-scores: 0.77–0.85; **Fig. 4B**). By day 25 post-IRI the distinction between and ischemic kidneys was very much reduced (F1-score: 0.58; **Fig. 4B**). These changing properties were also reflected in the spatial separation between contralateral control and ischemic tubules in **Fig. 4C–E**, but the separation of clusters was not visible at day 25 post-IRI (**Fig. 4F**). We characterized the structural changes imposed by IRI in contralateral control and ischemic kidneys over time to better understand the underlying biological manifestations of AKI. On day 1 post-IRI, there were significant difference in the microvasculature protein (96.4±25.6mg/mL *versus* 113.0±29.7mg/mL for contralateral control *versus* IRI, respectively; *p*≤0.001; *d*=0.61–0.67; **Fig. 4J**), possibly reflective of vascular congestion. On day 2 post-IRI, lipid droplet concentration was significantly lower in ischemic relative to contralateral control kidneys (92.8±22.7mg/mL *versus* 76.8±25.8mg/mL; *p*<0.001), whereas brush border lipid concentration was significantly greater in ischemic kidneys ([median±SEM] 30.7±0.4mg/mL *versus* 35.7±0.4mg/mL; *p*<0.001; **Fig. 4K**), however, effect size calculations revealed no effect (*d*=0.02) at day 2 post-IRI (**Fig. 4K**). On day 5 post-IRI, microvasculature protein concentration was also significantly higher in ischemic kidneys (89.0±23.3mg/mL *versus* 110.1±28.0mg/mL for contralateral control *versus* IRI, respectively; *p*<0.001; *d*=0.84; **Fig. 4L**), while lipid droplet size was significantly (but with small effect), elevated in ischemic kidneys (*p*<0.001; *d*=0.26; **Fig. 4L**).

**Figure 4.**
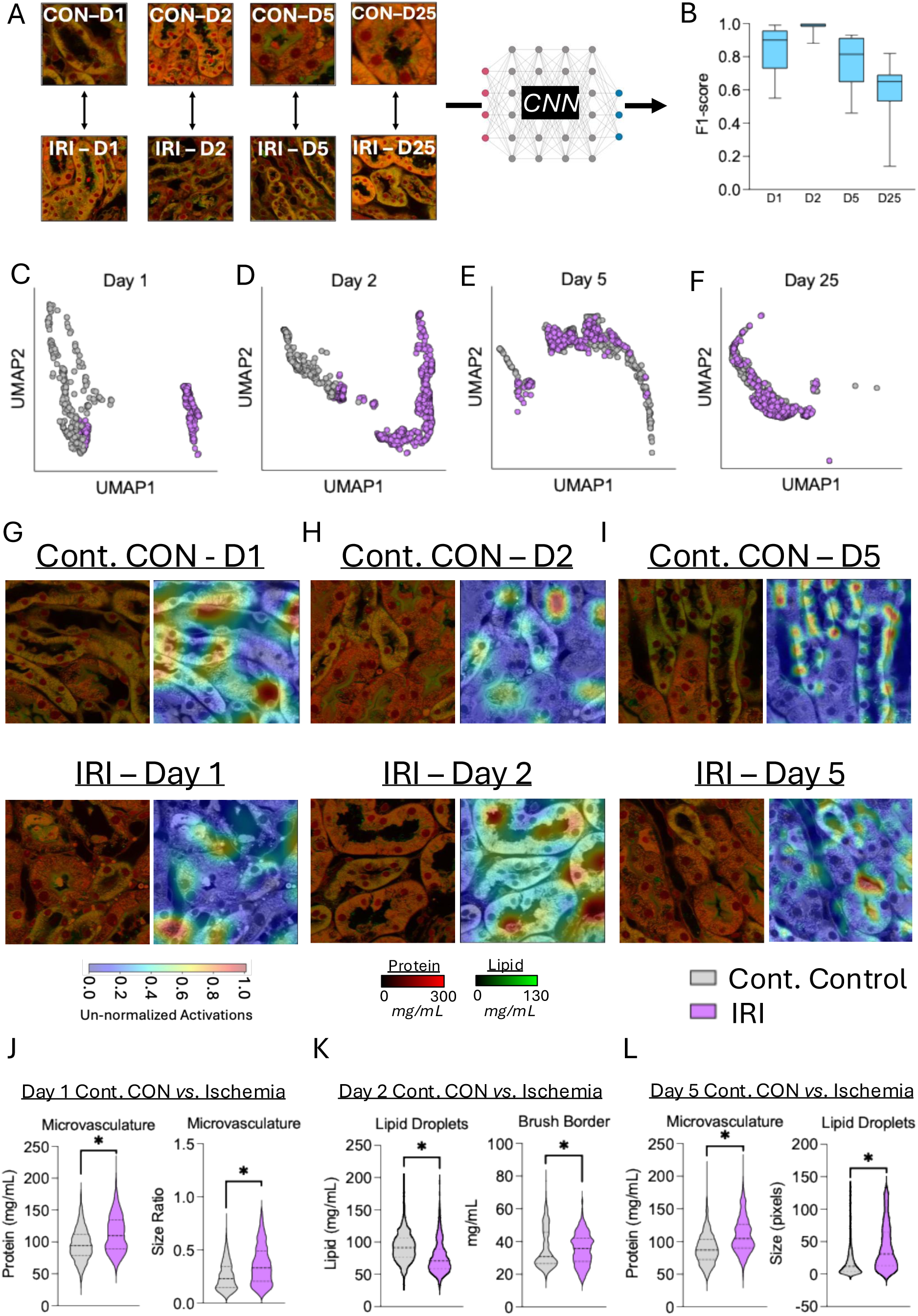
Exploration of the time-course effects of kidney ischemic reperfusion injury by NoRI. A, Representative NoRI images from each time point following ischemic reperfusion injury in both contralateral control (CON) and ischemic (IRI) kidneys input into ResNet50. B, Box-whisker plots of F1-scores for prediction of contralateral control vs IRI kidneys at each time point post-injury by ResNet50. Boxes represent median with 25^th^/75^th^ percentiles. Whiskers represent min-to-max values. C, UMAP plot of Cont. CON (contralateral control) (Grey) and IRI (Purple) tubules at day 1 post-injury predicted by ResNet50. Each datapoint represents an individual tubule. D, UMAP plot of Cont. CON (contralateral control) (Grey) and IRI (Purple) tubules at day 2 post-injury predicted by ResNet50. Each datapoint represents an individual tubule. E, UMAP plot of Cont. CON (contralateral control) (Grey) and IRI (Purple) tubules at day 5 post-injury predicted by ResNet50. Each datapoint represents an individual tubule. F, UMAP plot of Cont. CON (contralateral control) (Grey) and IRI (Purple) tubules at day 25 post-injury predicted by ResNet50. Each datapoint represents an individual tubule. G, Representative NoRI images from Cont. CON (contralateral control) and IRI kidneys at day 1 post-injury with adjacent heatmap of features defining CON from IRI identified by ResNet50. H, Representative NoRI images from Cont. CON (contralateral control) and IRI kidneys at day 2 post-injury with adjacent heatmap of features defining CON from IRI identified by ResNet50. I, Representative NoRI images from Cont. CON (contralateral control) and IRI kidneys at day 5 post-injury with adjacent heatmap of features defining Cont. CON (contralateral control) from IRI identified by ResNet50. J, Violin plots of microvasculature protein content and size measured by NoRI between Cont. CON (contralateral control) and IRI on day 1 post-injury. Violins are presented with median (dashed line) and 25^th^/75^th^ percentiles (dotted lines). K, Violin plots of lipid droplet concentration and brush border lipid concentration measured by NoRI between Cont. CON (contralateral control) and IRI on day 2 post-injury. Violins are presented with median (dashed line) and 25^th^/75^th^ percentiles (dotted lines). L, Violin plots of microvasculature protein concentration and lipid droplet size measured by NoRI between Cont. CON (contralateral control) and IRI on day 5 post-injury. Violins are presented with median (dashed line) and 25^th^/75^th^ percentiles (dotted lines). *Data in Panels J–L were analysed by Kolmogorov-Smirnov tests*.

We anticipated using the contralateral (unligated) kidney as an internal control for the IRI experiment. However, we were aware that the undisturbed contralateral kidney might have suffered an added burden for having to both undertake the function of two kidneys, as well as being exposed to a post-ischemic milieu of stress and damage from response molecules released by the ligated kidney. Thus, we next turned our attention to detecting the effects of IRI on the unligated contralateral control kidney and in ischemic kidney in isolation, over time (**Fig. 5A**). There were, in fact, significant changes over time within both the contralateral control and ischemic kidneys over time (F1-score: 0.77–0.99; **Fig. 5B**), suggesting the contralateral control kidney also exhibits substantial temporal responses post-IRI. Specifically, distinct clustering between days 1 *versus* 2 and days 5 *versus* 25 was apparent in the contralateral control kidney data (**Fig. 5C and E**). Indeed, this separation in the contralateral control kidney UMAP clustering was similar to the distinct clustering found between timepoint post-IRI in the ischemic kidneys (**Fig. 5F–H**). Using ResNet50 CNN analyses on the quantitative, undisrupted NoRI images of kidneys we could easily detect changes in both contralateral control and ischemic kidneys post-IRI (*representative images presented in* **Fig. 5I**). We found the concentration of protein within the tubule lumen (representative of a pathological ‘protein plug’) increased from 30.8±19.3mg/mL to 39.9±25.0mg/mL, in contralateral control kidneys, and decreased from 37.1±24.2mg/mL to 34.7±22.8mg/mL from day 1 to day 25 post-IRI in ischemic kidneys (**Fig. 5J**), but was not entirely absent by day 25 post-IRI. Lipid droplet concentrations also increased from 88.0±31.2mg/mL to 104.3±27.7mg/mL, in contralateral control kidneys, and from 93.8±32.9mg/mL to 94.9±26.4mg/mL from day 1 to day 25 post-IRI in ischemic kidneys (**Fig. 5K**). Finally, we found brush border lipids also increased from 35.4±11.2mg/mL to 43.7±11.0mg/mL, in contralateral control kidneys, and from 34.4±11.3mg/mL to 37.6±9.7mg/mL from day 1 to day 25 post-IRI in ischemic kidneys (**Fig. 5M**). However, the comparative lack of brush borders at early time points (pre-day 5 post-IRI) may have caused a skewing of these analyses towards ‘less-perturbed’ regions that still had brush border present for analysis relative to tubules that entirely lost their brush border and so would be precluded from this analysis. All these features displayed highly significant time effects from day 1 to 25 post-IRI (*two-way ANOVA time effect*: *p*≤0.0001; **Fig. 5J–M**). Not surprisingly, there were significant interaction effects (all *p*≤0.0001) between contralateral control and ischemic kidneys at each timepoint for each of the four anatomical regions quantified in Panels J–M, showing differential effects of ischemia on contralateral control and ischemic kidneys.

**Figure 5.**
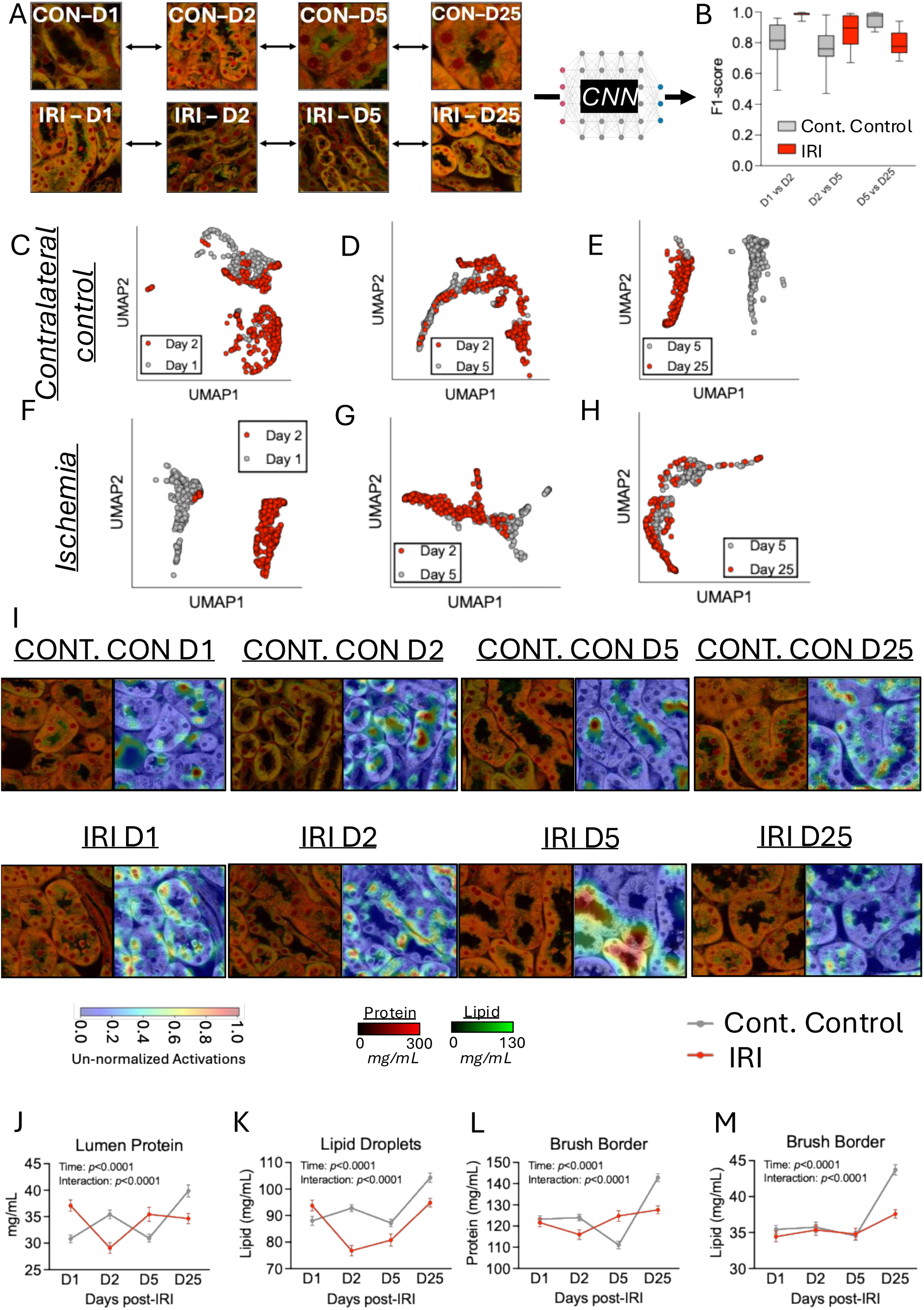
Time course analysis of the effects of ischemic reperfusion injury within contralateral control and IRI kidneys. A, Representative NoRI images from each time point following ischemic reperfusion injury in both contralateral control (CON) and ischemic (IRI) kidneys input into ResNet50. B, Box-whisker plots of F1-scores for prediction of the time point post-injury for Cont. Control (contralateral control) and IRI by ResNet50. Boxes represent median with 25^th^/75^th^ percentiles. Whiskers represent min-to-max values. C, UMAP plot of day 1 (grey) and day 2 (red) tubules for CON kidneys post-injury predicted by ResNet50. Each datapoint represents an individual tubule. D, UMAP plot of day 2 (grey) and day 5 (red) tubules for CON kidneys post-injury predicted by ResNet50. Each datapoint represents an individual tubule. E, UMAP plot of day 5 (grey) and day 25 (red) tubules for CON kidneys post-injury predicted by ResNet50. Each datapoint represents an individual tubule. F, UMAP plot of day 1 (grey) and day 2 (red) tubules for IRI kidneys post-injury predicted by ResNet50. Each datapoint represents an individual tubule. G, UMAP plot of day 2 (grey) and day 5 (red) tubules for IRI kidneys post-injury predicted by ResNet50. Each datapoint represents an individual tubule. H, UMAP plot of day 5 (grey) and day 25 (red) tubules for IRI kidneys post-injury predicted by ResNet50. Each datapoint represents an individual tubule. I, Representative NoRI images from CONT. CON (contralateral control) and IRI kidneys at days 1, 2, 5, and 25 post-injury with adjacent heatmap of features defining each time point, within CON and IRI kidneys identified by ResNet50. J, Line graph of lumen protein concentration measured by NoRI at each time point post-injury for CONT. CON (contralateral control) (grey) and IRI (red). The mean value for each group at a given time point post-injury is presented, with error bars representing SEM. K, Line graph of lipid droplet concentration measured by NoRI at each time point post-injury for CONT. CON (contralateral control) (grey) and IRI (red). The mean value for each group at a given time point post-injury is presented, with error bars representing SEM. L, Line graph of brush border protein concentration measured by NoRI at each time point post-injury for CONT. CON (contralateral control) (grey) and IRI (red). The mean value for each group at a given time point post-injury is presented, with error bars representing SEM. M, Line graph of brush border lipid concentration measured by NoRI at each time point post-injury for CONT. CON (contralateral control) (grey) and IRI (red). The mean value for each group at a given time point post-injury is presented, with error bars representing SEM. *Data in Panels J-M were analysed by two-way repeated measured ANOVA with Bonferroni post-hoc corrections applied*.

To understand the power of the quantitative nature of NoRI data when combined with CNNs we examined kidneys from day 2 post-IRI for deeper analysis. We found a considerable improvement in F1-scores (as a measure of accuracy of predicting ischemic from contralateral control kidney images) when we utilized both protein and lipid versus just protein alone (F1=0.84 *versus* 0.99, respectively). This gave us our first sense of the importance of the lipid information, which cannot be studied using conventional histology techniques since lipids are extracted during the embedding procedure. We expanded these analyses by asking how important the absolute quantitative information obtained by NoRI was, compared with less quantitative measurements, for the simple task of distinguishing contralateral control from ischemic kidneys at day 2 post-IRI. To make this comparison we generated non-quantitative NoRI images where samples were normalized on a per-image basis, and protein and lipid were also normalized in isolation from one another (similar to intensity-based histogram adjustments in conventional histology). As expected, there was a reduction in F1-scores from quantitative to non-quantitative data, with a further drop between non-quantitative data when only protein is used compared to protein and lipid together (**Supplementary Fig. 13**).

## Discussion

NoRI is a novel microscopy technique that provides a highly quantitative measure of both protein and lipid biomass concentrations at high spatial resolution. As such it represents a potentially powerful new tool for detecting subtle tissue injury or physiological state change. This is the *a priori* argument, but it is only in the setting of real pathology that one can assess the value of any new histopathological method. Hence, our goal in this study was to test NoRI on a realistic problem of tissue physiology; we chose ischemic reperfusion injury in the kidney. We found that in the setting of fully quantitative, spatial information where there was no disturbance to tissue architecture, as provided by NoRI, we were able to characterize, and for the first time, quantitate known features of AKI (e.g., luminal protein plugs and brush border loss), and discover novel features of kidney damage and reparative processes that would not have been possible previously (e.g., lipid droplet concentration heterogeneity, brush border lipid concentrations). Further, we found that precisely because NoRI images are fully quantitative, normalised to a reference calibration on each day of use, and do not require tissue processing, and hence do not cause disruption of tissue architecture, they are fully reproducible. As such, they are the ideal image data for analysis by neural networks. This combination of advanced, unbiased analysis methods, such as with NoRI images, can provide a unique platform to explore and identify previously unknown biological phenomena in the setting of complex tissues, and at the same time to compare the effectiveness of different methods to do so.

The standard protocols in histopathology involve fixation, dehydration, generally paraffin embedding, and sectioning. These processes extract lipids, which represent a considerable portion of mammalian cellular biomass. Routine clinical pathology uses an array of techniques, such as stains like hematoxylin and eosin, most recently complemented by immunohistochemistry, usually on microtome-cut, paraffin-embedded tissue slices. Owing to such harsh treatments required by the needs to preserve some tissue architecture and the hardening of tissue that can then be sectioned to thicknesses of less than 10µm, there is only partial preservation of the *in vivo* architecture. These sequential preparation steps are also time-intensive (as much as several days), and the indirect staining/probing methods exhibit day-to-day variability. As such, while relatively inexpensive, the traditional methods provide an incomplete and imperfect record of tissue architecture. Conversely, the imaging and analytical work, built upon our initial work using NoRI(3), overcomes most of the traditional constraints. Indeed, NoRI is performed in thick (100–500µm, and perhaps up to 1mm) tissue samples that are optically sectioned, rather than mechanically sliced (with an axial resolution of approximately 1.9µm(3)). This allows 3-dimensional high-resolution imaging of cells and organelles and notably does not require tissue processing. It can be performed on formaldehyde-fixed, unfixed or living material(3, 4). It can provide a fully quantitative readout of protein and lipid concentrations, and, by varying the wavelength, can distinguish chemical differences, such as the degree of saturation of lipids and the quantitative levels of cholesterol. With proper protocols in place (i.e., where appropriate machine learning models have been trained prior), imaging and analysis can be performed in <1 hour of tissue presentation. Furthermore, NoRI is fully compatible with immunofluorescence.

As discussed above, NoRI images can be processed quickly (<1 hour) compared to conventional histology (hours-to-days). Once acquired, we have developed protocols for automatically segmenting kidney tubules and substructures (nuclei, nucleoli, brush border, lumen) using two open-access methods—SAM and YOLO v.8—that performed with high precision when compared with laboriously obtained hand-segmented reference data. This unexpected facility can largely be explained by the reproducibility and quantitative nature of NoRI data. Spectrophotometric detection of optimally sectioned material is computationally unmixed to offset any impact of light scattering in complex samples. After trying many computation methods, we settled on an open-access convolutional neural network (ResNet50); it provided a powerful and fast means of high-dimensional analysis of NoRI image. Indeed, we have shown how, with very little input training data (as few as 2–3 input images), we are able to identify three distinct tubule types, anatomical location, biological sex, and disease status of tubules with high confidence. While cytoplasmic protein and lipid concentrations were sufficient to delineate proximal from distal tubules with reasonable confidence, the use of protein, lipid, and geometric data in the ResNet50 pipeline was able to determine different tubule types with near-perfect (against the gold standard of human annotation) certainty. We further discovered the context-specific importance of lipid and protein concentrations during this process. For example, we found that lipids were the most important feature to distinguish contralateral control from ischemic kidneys at day 2 post-IRI, suggesting lipids are central to the early pathology of acute tubular injury. Conversely, protein concentrations were a stronger determinant of predicting the sex of the kidney being analysed, indicating lipid handling within the kidney is more congruent between male and female tubules than is protein. These results importantly show the value of NoRI quantification of both proteins and lipids.

Another important characteristic of the quantitative protein and lipid data obtained by NoRI, coupled with the preservation of tissue architecture due to the lack of tissue processing required for NoRI, was the ability to detect small structures of importance in these images. This was made most clear when we applied the same machine learning analysis approaches to both NoRI and H&E sections from the same male and female mice for direct comparison. The value of this finding is in support of the lack of sample processing, and thereby greater preservation of *in situ* tissue architecture with NoRI imaging compared to conventional histology techniques. Using NoRI image data, the neural network models we developed were able to identify, consistently, small and refined structures such as lipid droplets to distinguish males from females. Comparatively, H&E images, analysed using the same approaches, were unable to identify the sex of the kidney from quantitation of such small structures, and instead only identified larger structures, such as tubules, as distinguishing features of male and female kidneys. This side-by-side comparison of NoRI *versus* H&E in the same specimens highlighted that, when combining NoRI with these advanced analysis techniques, we were uniquely positioned to make striking insights into healthy and diseased mouse kidneys.

### The utility of NoRI in histology and pathology

We chose the kidney as a model system for testing the effectiveness of NoRI, combined with neural networks/machine learning methods for the analyses of tissue histology and/or pathology. The kidney is comprised of highly repeated structures (tubules), distinct anatomical characteristics (cortex/medulla), distinct sub-structures (cytoplasm, brush border, lumens), and a variety of cell types. With these attributes in mind, we addressed three biological questions: the extent and nature of the kidney’s anatomical variability, its sexual dimorphism, and anatomical changes in the response to ischemia-reperfusion injury (as a model of Acute Kidney Injury [AKI]).

Using NoRI alone we found distinct clustering of cell types solely by their concentration of protein and lipid. This was confirmed by the unique expression of transporters and channels revealed by specific antibodies. The distal tubules, proximal tubules, and collecting ducts are epithelial cells expressing different membrane antigens. Yet, unexpectedly, the cytoplasm of proximal tubules was significantly higher in total protein on average by 24mg/mL and lower in lipid on average by 10mgmL. Collecting ducts were lower in both protein and lipid than either proximal or distal tubules by 32-56mg/mL and 12-22mg/mL, on average, respectively. What was also notable from these results was how tightly-regulated the cytoplasmic concentrations of proteins and lipids were within tubule types. The protein concentrations varied on average 17.2mg/mL (9.5%), while the lipid concentrations varied 9.9mg/mL (17.2%). We also found that the nuclear protein content varied between tubule types, with proximal tubules having ∼30mg/mL higher protein concentrations than distal tubules or collecting ducts. Before NoRI measurements, such information on the distinctive lipid and protein concentrations in the cytoplasm of kidney tubules has not to our knowledge been previously reported. Differential cytoplasmic lipid content between tubule types may be related to metabolic differences, with proximal tubules biased towards lipid metabolism, evidenced by a greater abundance of PPARα/γ—involved in lipid metabolism—while distal tubules are more glycolytic in nature(23), and therefore less reliant on an intracellular lipid pool. The lower cytoplasmic protein density of proximal tubules may be due to their higher lipid content and enhanced rate of lipid oxidation/turnover. Such observations suggest that quantitative features of protein and lipid density may reflect physiological conditions and be more informative than are morphological phenotypes.

Having established that we could determine tubule identify with high confidence, we investigated the anatomical variation of the kidney protein and lipid as related to anatomy. We first examined nephrons in three adjacent anatomical regions—Rostral pole, Midpole, and Caudal pole portions—from cortex to medulla.

These three regions were easily distinguishable by NoRI, as exemplified by protein concentration difference in the microvasculature, which ranged 77.2—100.2mg/mL, and was almost 30mg/mL higher in Caudal relative to Rostral or Midpole regions. Likewise, lipid droplet concentrations were considerably different between regions, being highest in Caudal pole sections, whereas lipid droplet size was nearly 3-fold greater in Midpole regions relative to Rostral and Caudal pole sections. These results represent an intriguing insight into the kidney microvasculature. Prior work utilising super-resolution ultrasound imaging in Zucker rats identified the Midpole as the region most dense in microvasculature(24), while we found protein concentration was greatest in the Caudal pole region. This may, of course, represent a species difference that remains to be determined, but may also represent a subtle difference in the microvasculature itself. For instance, should microvasculature be most dense in the Midpole region—as suggested by McDermott*, et al.* (24)—the endothelial cells themselves may have differential cytoplasmic protein concentrations, reflecting the higher Caudal protein concentrations relative to Midpole or Rostral pole regions in the present study.

It is well-known that mouse kidneys exhibit some degree of sexual dimorphism, with larger kidneys in males, reflecting differential water and sodium transporter expression(25-27). Human male and female kidneys show different rates of decline in function with age(27), differential endothelial integrity, differential metabolic profiles, and divergent responses to ischemic-reperfusion injury (*summarized in* Steiger*, et al.* (28)). These differences are most pronounced in the cortex and outer stripe of the medulla(29). Consistent with these findings, we were easily able to distinguish male from female kidneys at a structural level with high reproducibility in the NoRI images. On the compositional level, we found cytoplasmic protein content was significantly higher in proximal tubules from female mice (256.9±6.9mg/mL) *versus* males (245.5±3.2mg/mL). By contrast, cytoplasmic lipid was more concentrated across all tubules in female mice (males: 50.2±2.0mg/mL *versus* females: 57.1±2.5mg/mL), which could be consistent with the higher resilience to lipid accumulation in female relative to male kidneys(30). This was also supported by another feature: our finding that lipid droplets were, on average, almost 7.2µm^3^ larger in female kidneys and brush border lipid concentration was 11.65mg/mL more concentrated, on average, in females compared to males. Furthermore, the protein content of the nucleus was also, on average 26.3mg/mL higher in female tubules. It is also known that female brush border transporter expression and function also differ from males, which may be partially androgen-related(30, 31). While the molecular basis underlying molecular differences in the protein content of the nucleus is unknown and the reasons for the much greater lipid density in female brush borders remain uncertain, these findings show that there are several sexual dimorphisms in murine kidneys, which NoRI has been able to provide the first quantitative readouts for. The power of such quantitative measurements by NoRI is significant. Further, we have also shown how, using NoRI coupled with neural network analyses, we can delineate these sexual dimorphisms in regions as small as 16.6µm^2^, with ≥85% certainty, emphasizing the pervasive and consistent characteristics of sexual dimorphism in the kidney With this morphological and structural consistency in mind, we applied NoRI to the ischemic-reperfusion injury model of kidney dysfunction and repair that resembles AKI in humans(22, 31, 32). We can replicate the natural event by transiently interrupting the blood supply to one kidney via clamping the renal artery leaving the contralateral kidney unperturbed directly. As discussed further below, we initially assumed the unperturbed kidney would act as a convenient internal contralateral control. The ensuing hypoxia, caused by a transient clamping of the renal artery leads to both acute functional and structural injury to the ligated kidney, and following 25 days from injury, the kidney recovers much of its function and repairs the damage. Histologically, the transient damage results in sporadic patches of injured tubules and necrosis, characterised by protein plugs in the luminal regions and loss of brush borders. During the recovery phase there are morphologically distinct recovering tubules, and seemingly spared healthy tubules(32). It is known that the relative abundance of these damaged and healthy regions depends on the ischemia duration, age, sex, and underlying disease(32). Ischemic injury is known to cause both endothelial and epithelial cell breakdown and swelling(22), which would likely explain the features identified in NoRI images. The accumulation of protein within the inter-tubule regions may perhaps also be reflective of an accumulation of leukocytes and fibroblasts around dead/dying tubules as part of the adaptive response to injury(22, 31, 32), especially in male kidneys(33). We also found day 2 post-injury was hallmarked by differences in lipid droplet concentrations (on average 16mg/mL higher in ischemic relative to contralateral control kidneys) in proximal tubules. Specifically, we observed a steady increase in both protein and lipid within brush border regions by day 25 post-injury relative to day 1 with an average increase of 8.3mg/mL and 3.1mg/mL protein, and 16.2mg/mL and 1.1mg/mL lipid concentration in contralateral and ischemic kidneys, respectively. This likely represents the re-establishment of the brush border (more marked in contralateral control than ischemic kidneys).

The large changes in lipid amount and distribution suggest that metabolic changes are a central feature of ischemic reperfusion injury. For example, as discussed above, we identified lipid droplet concentrations, and heterogeneity in droplet concentration, as well as brush border lipid concentrations as major determinants of the injury and repair process during this model of AKI. Kidney proximal tubules are heavily reliant on fatty acid oxidation as a primary source of energy production. Defective lipid handling is a hallmark of renal dysfunction and fibrosis(34, 35). It is, therefore, interesting to quantitatively capture distinct changes in lipid droplet concentrations following ischemic injury in both contralateral control and ischemic kidneys, within their spatial context. These observations are greatly facilitated by NoRI given it provides the first quantitative readout of total lipids in AKI, as lipotoxicity is established as a major hallmark of AKI, and a focus of therapeutic developments(35). Lipid droplets are a heterogeneous population of phospholipid monolayer-bound organelles that store neutral lipids, usually for later use by metabolic systems(36, 37). When the lipid supply outweighs utilization, a lipid-storage response is initiated(38). Therefore, it could be expected that ischemic injury, which causes substantial impairments to beta-oxidation capacity in tubules and a switch to anaerobic metabolism(38, 39) would lead to substantial lipid accumulation. The transport of lipid from the blood might be unlinked from the utilization of lipid, which requires oxidative respiration, which has been inhibited by hypoxia. This imbalance would be expected to lead to an accumulation of lipid droplets during transient hypoxia. Though this might seem reasonable, we in fact observe just the opposite, a lower lipid droplet concentration in ischemic kidneys relative to contralateral control kidneys, notably at day 2 post-IRI (92.8±22.7 *versus* 76.8±25.8mg/mL for contralateral control and ischemic kidneys, respectively). This trend persisted to day 25 with lipid droplet concentrations being ∼6.5 and 9.4mg/mL lower in ischemic relative to contralateral control kidneys. While the density of lipid droplets may change in response to changes in their biophysical properties and constituent lipids (triglycerides, cholesterol esters, fatty acids), the presence of coalesced regions of lipids following ischemic-reperfusion injury may not represent physiological lipid droplets *per se* and instead may represent pools of lipids within the debris of dead/dying cells. Lipid-droplet membrane protein probing by immunofluorescence may clarify this uncertainty in future work. Thus, if this undirected accumulation of lipids occurs from the remnant lipids from within the dead cells’ cytoplasm, then their density could feasibly be less than organelle-bound lipids. Indeed, the observation of significantly large lipid droplets by day 5 post-injury could support this suggestion.

An important contribution made by NoRI in the context of the reperfusion-injury model of acute kidney injury used here was the profound influence the injury imparted on the contralateral control kidney over time. Our quantitative histopathology investigations of the responses over time in the contralateral control kidneys identified many features of dysfunction and damage. Notably, we were able to distinctly separate individual time points between days 1 to 25 post-IRI in the contralateral control group, specifically, by performing neural network analysis on NoRI images. Specifically, these models highlighted a gradual increase in luminal protein (i.e., protein plugs) from days 1 to 25, as well as a gradual increase in lipid droplet concentrations, similar to what was observed in the ischemic kidneys. Furthermore, by day 25 post-IRI there was a dramatic increase in brush border protein and lipid concentrations in the contralateral control kidneys, relative to days 1–5 post-IRI, suggestive of a re-establishment of the brush border that was previously reduced in size and density in the contralateral control kidneys. Our observations support prior work that show the structural, transcriptional, and functional dysfunction in the contralateral control kidneys of this model of AKI, but crucially add additional, quantitative data to this body of work.

The dysfunction in the contralateral control kidneys can be segregated into short-term (hours/days), medium-term (days/weeks), and long-term (weeks/months) post-injury. In the short-term, contralateral control kidneys exhibit strong transcriptional shifts due to oxidative stress, increased inflammatory cytokines, and metabolic dysfunction(40, 41). Indeed, the contralateral kidney is exposed to the cytokine storm and oxidative stress produced by the ischemic-damaged kidney, which can influence the healthy kidney that is responsible for a greater proportion of the biological function of two kidneys. In this context, we found a notable change in lipid droplet concentrations from ∼90mg/mL on day 1 post-IRI to ∼105mg/mL by day 25. This provides a quantitative measurement supporting and expanding upon previous reports of downregulated lipid metabolism pathways in the contralateral control kidney within 24-hours of injury(42). Our results represent the first highly sensitive, quantitative measurement of lipid-related effects of ischemic-reperfusion injury, to our knowledge. In the weeks following injury, leukocytes, including activated T-cells, infiltrate both kidneys. This can represent both part of the damage-induced inflammatory program as well as the structural remodelling—in concert with fibroblasts—that occurs within the contralateral kidney as it compensates for the loss of the damaged kidney(43). In this respect, our analyses identified several features of both tubule damage/stress and structural remodelling within the contralateral kidney. These were hallmarked by the gradual increase in luminal protein from ∼32mg/mL on day-1 to ∼40mg/mL on day-25 post-injury. Likewise, brush borders showed quantitative evidence of either re-establishment post-injury, or increased abundance and/or density of structures by day-25 post-injury, evidenced by a dramatic increase in protein (from ∼125mg/mL to ∼140mg/mL) and lipid (from ∼36mg/mL to ∼44mg/mL) concentrations at day-25 relative to days 1–5 post-injury. Such quantitative tracking of the micro-anatomy of the kidney in the context of AKI constitutes an exciting development of both known features of kidney damage and repair as well as possibly presenting novel features of injury and repair. These data represent initial insights into the power of NoRI as a tool for making novel biological insights into pathological states, as well as being a useful diagnostic tool in histopathology.

## Conclusion

We present this first exploration of the potential of NoRI—a fully quantitative normalized form of stimulated Raman spectroscopy that measures protein and lipid content in biological samples— combined with neural network analysis as a tool for feature identification and diagnostics in histopathology. We have shown, using murine kidney as a candidate organ for these investigations, that NoRI is fully capable of analysing complex tissue samples in a way that could be considered superior to conventional histology methods, owing to its comparatively fast turnaround time from sample harvesting to imaging, its ability to analyze samples without the need for destructive or damaging sample processing, and its ability to fully quantify protein and lipids, label-free, at high spatial resolution. Moreover, we have shown how this form of microscopy, due to its absolute (as opposed to relative) quantitation is highly appropriate for analysis by neural networks for feature discovery. Because it is a spectrophotometric absolute measurement, it releases the power of machine learning for unbiased feature discovery, suggesting that it may have strong diagnostic potential in physiology and disease. Finally, due to the reproducibility (both between animals and within tubule types) and the quantitative nature of these data, these advanced analysis approaches require a relatively low level of input data for feature discovery, such that fine-tuned models can be easily derived and applied to related samples. Thus, NoRI could be applied to a vast array of biological questions and disease states to make both novel biological discoveries and facilitate disease diagnostics in the clinical context. Of these questions, both discovery of novel foci, and the early diagnosis of cancers are fundamental questions that NoRI is well-placed to make advancements in. Likewise, the application of NoRI to the ageing process could add fundamental insights into a poorly understood, biologically relevant phenomenon, especially given the accumulation of lipids in many organs with age, and the fact that senescent cells are characterised by cell swelling and cytoplasmic mass-density uncoupling.

## Author Contributions

*Research design and conception*: S.O., W.V.T., T.I., J.V.B., L.P., and M.K.

*Primary data collection*: S.O., W.V.T., T.I., A.T.

*Data analysis and interpretation*: S.O., W.V.T., M.D., K.P., T.I., A.T., J.V.B., R.K., S.F.N., L.P., and M.K.

*Manuscript writing*: S.O., W.V.T., M.D., and M.K.

## Acknowledgements

## Acknowledgements

We would like to thank Dr. ChangHee Lee for his helpful assistance in providing the FiJi script for NoRI channel assembly. We would also like to thank Dr. Xili Liu, Dr. Avik Mukherjee, and Ang Li for their helpful assistance in maintaining the NoRI microscope.

## Data availability

All raw code is available at: https://github.com/kirschnerlab/NoRI

https://github.com/HMS-IAC/nori

https://github.com/MDyakova/NORI_image_segmentation

https://github.com/MDyakova/NORI_feature_extraction

https://hms-iac.github.io/NoRI/

All raw image data are available at Zenodo: https://doi.org/10.5281/zenodo.17329675.

