## Supplementary Figures for "Normalized Raman Imaging for Studies of Tissue Physiology of the Kidney"

**Supp. Fig. 1**

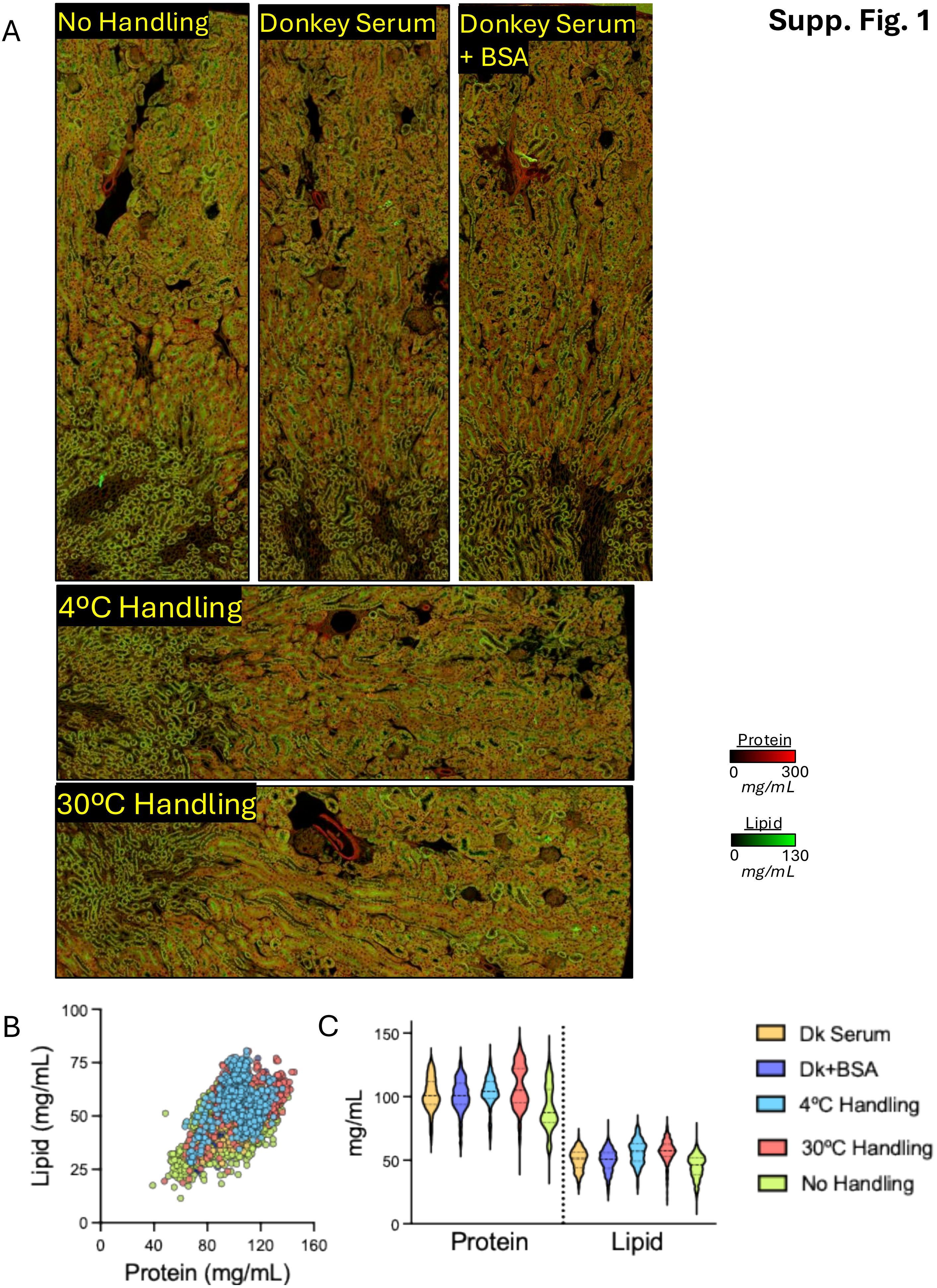
A
No Handling
Donkey Serum
Donkey Serum + BSA
4ºC Handling
30ºC Handling
Protein
0 300
mg/mL
Lipid
0 130
mg/mL
B
Lipid (mg/mL)
Protein (mg/mL)
0
25
50
75
100
0
40
80
120
160
C
mg/mL
0
50
100
150
Protein
Lipid
Dk Serum
Dk+BSA
4ºC Handling
30ºC Handling
No Handling

**Supp. Fig. 2**

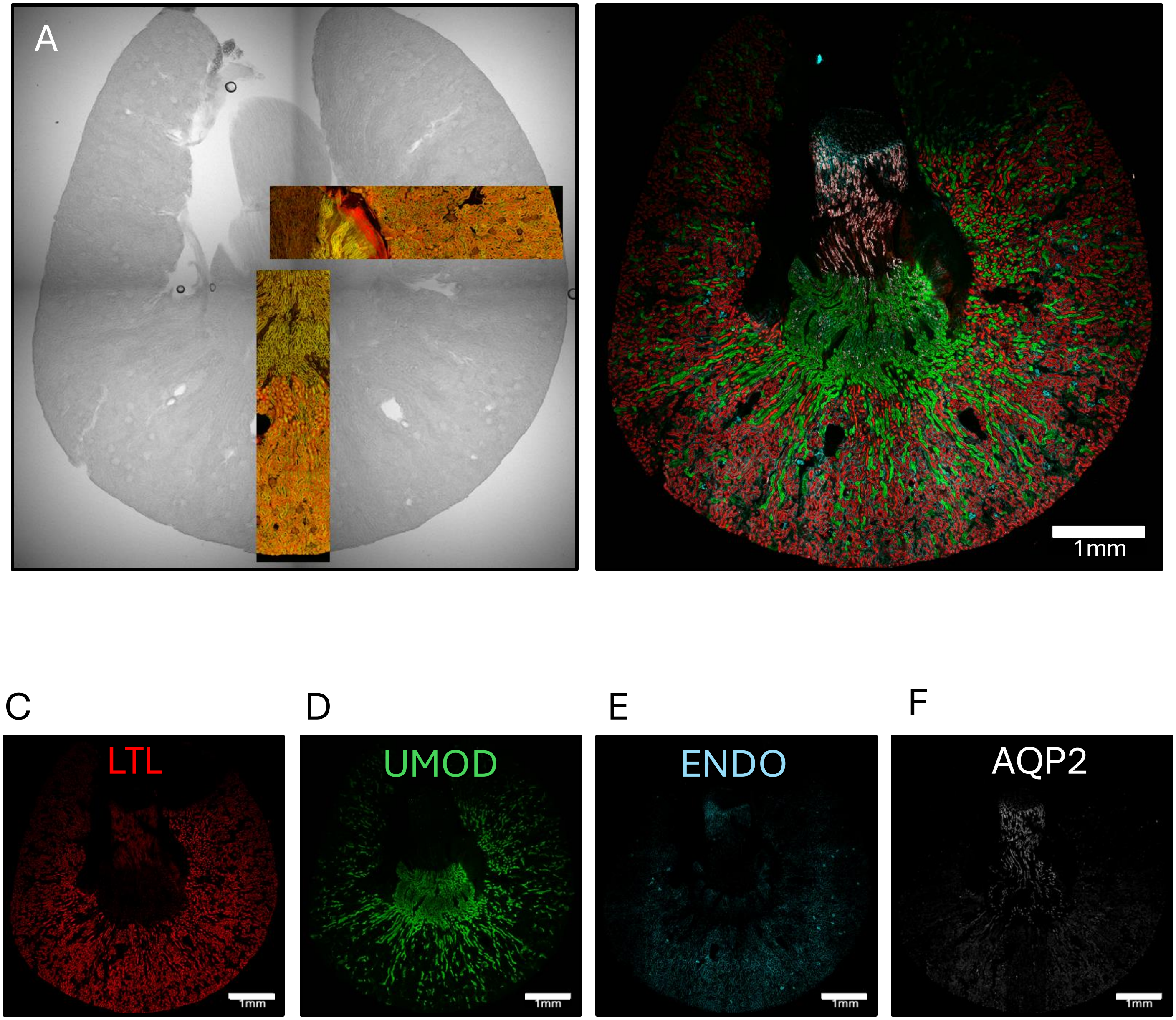

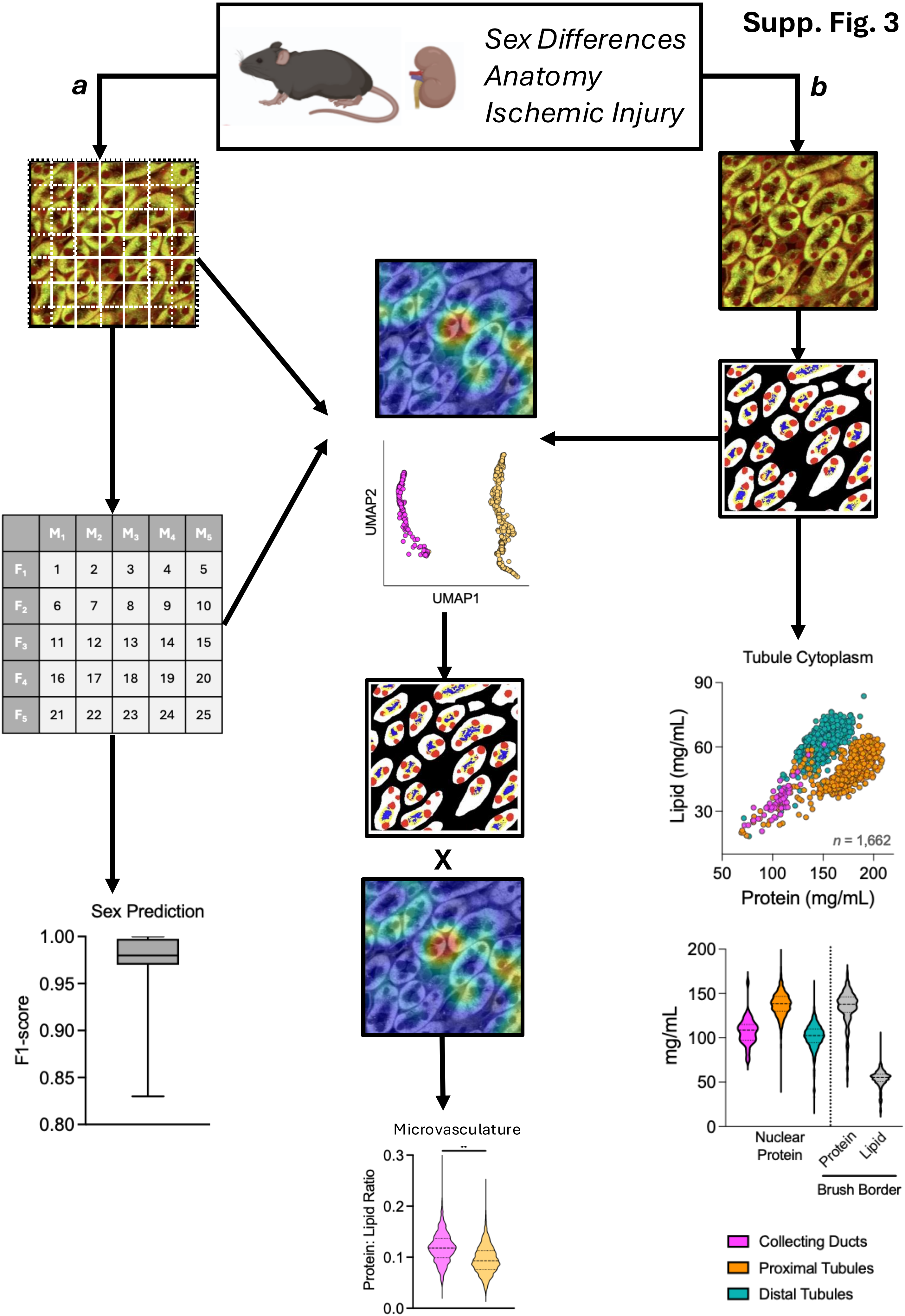
Supp. Fig. 3
Sex Differences
Anatomy
Ischemic Injury
a
b
M1
M2
M3
M4
M5
F1
F2
F3
F4
F5
Sex Prediction
F1-score
1.00
0.95
0.90
0.85
0.80
UMAP2
UMAP1
X
Microvasculature
Protein: Lipid Ratio
0.3
0.2
0.1
0.0
**
Tubule Cytoplasm
Lipid (mg/mL)
90
60
30
n = 1,662
50
100
150
200
Protein (mg/mL)
mg/mL
200
150
100
50
0
Nuclear
Protein
Protein
Lipid
Brush Border
Collecting Ducts
Proximal Tubules
Distal Tubules

**Supp. Fig. 4**

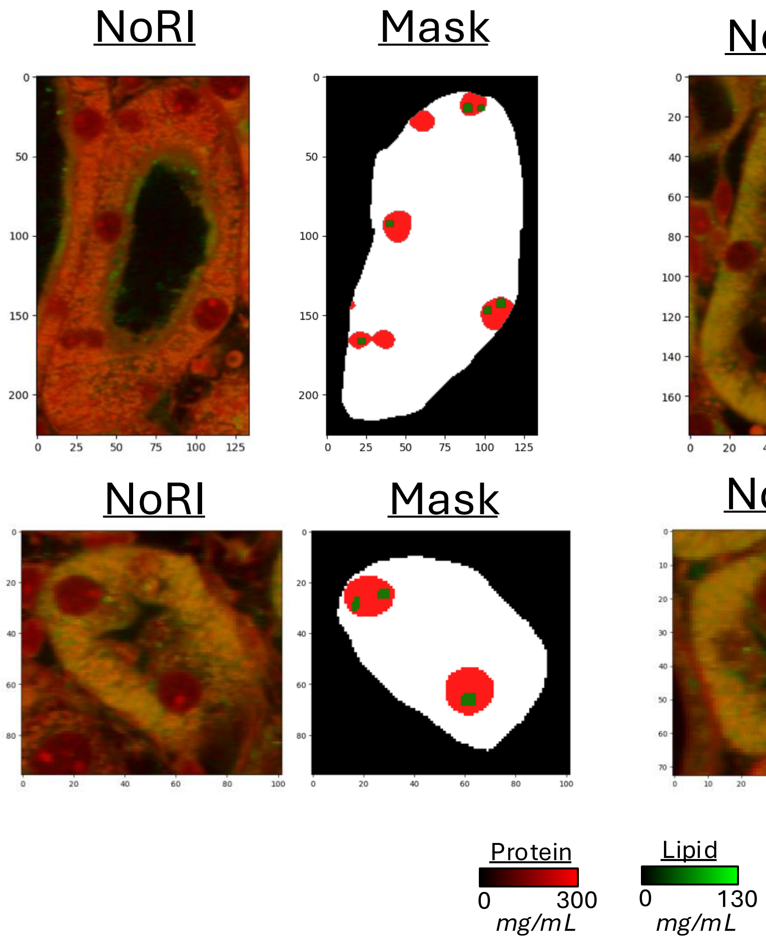

NoRI
Mask
NoRI
Mask
NoRI
Mask
NoRI
Mask
Protein
0
300
mg/mL
Lipid
0
130
mg/mL

**Supp. Fig. 5**

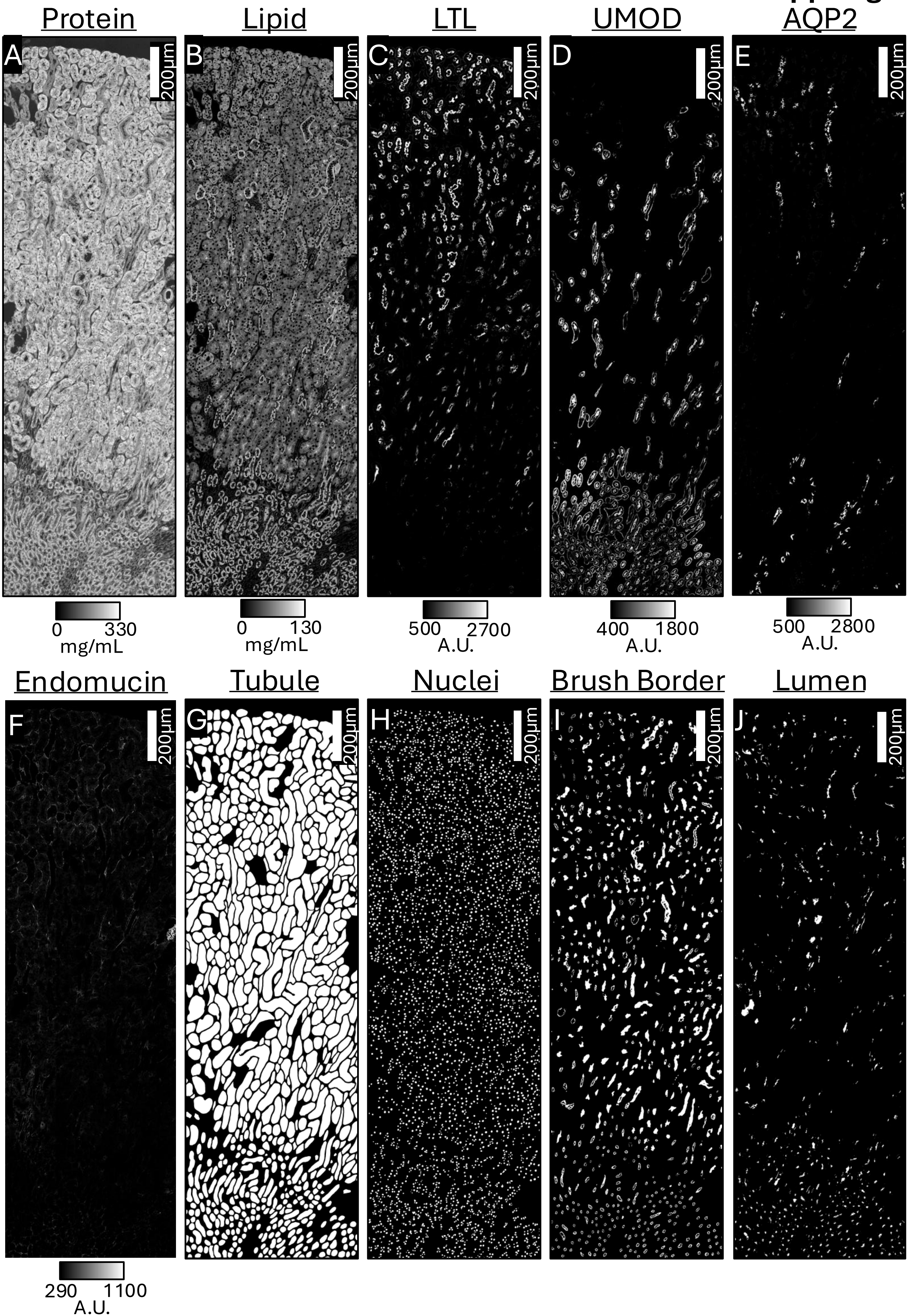
Protein
Lipid
LTL
UMOD
AQP2
A
B
C
D
E
200µm
0 330 mg/mL
0 130 mg/mL
500 2700 A.U.
400 1800 A.U.
500 2800 A.U.
Endomucin
Tubule
Nuclei
Brush Border
Lumen
F
G
H
I
J
290 1100 A.U.

**Supp. Fig. 6**

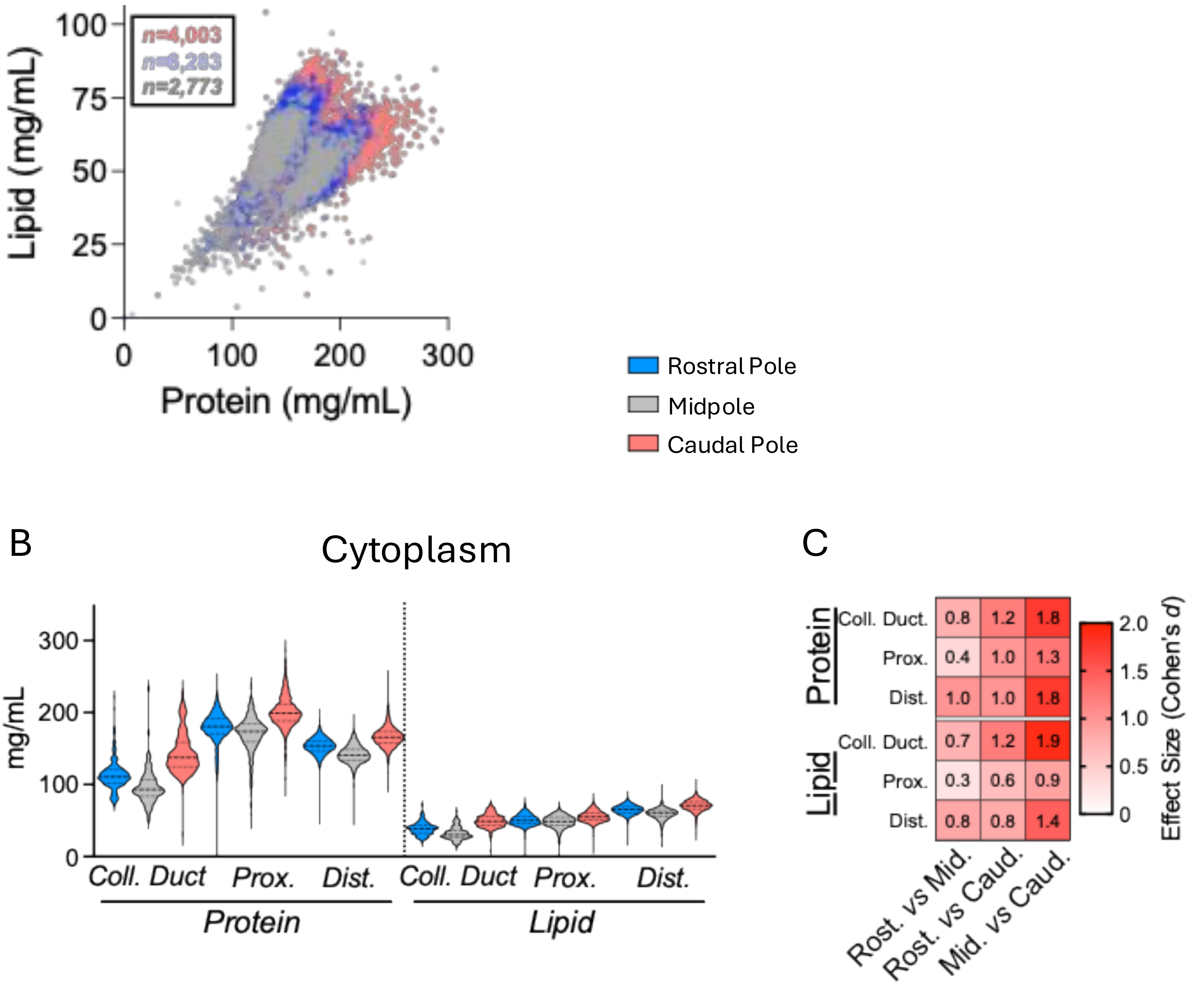
A
Lipid (mg/mL)
Protein (mg/mL)
n=4,003
n=6,283
n=2,773
0
25
50
75
100
0
100
200
300
Rostral Pole
Midpole
Caudal Pole
B
Cytoplasm
mg/mL
300
200
100
0
Coll. Duct
Prox.
Dist.
Coll. Duct
Prox.
Dist.
Protein
Lipid
C
Protein
Lipid
Coll. Duct.
Prox.
Dist.
Coll. Duct.
Prox.
Dist.
0.8 1.2 1.8
0.4 1.0 1.3
1.0 1.0 1.8
0.7 1.2 1.9
0.3 0.6 0.9
0.8 0.8 1.4
Rost. vs Mid.
Rost. vs Caud.
Mid. vs Caud.
Effect Size (Cohen's d)
2.0
1.5
1.0
0.5
0

**Supp. Fig. 7**

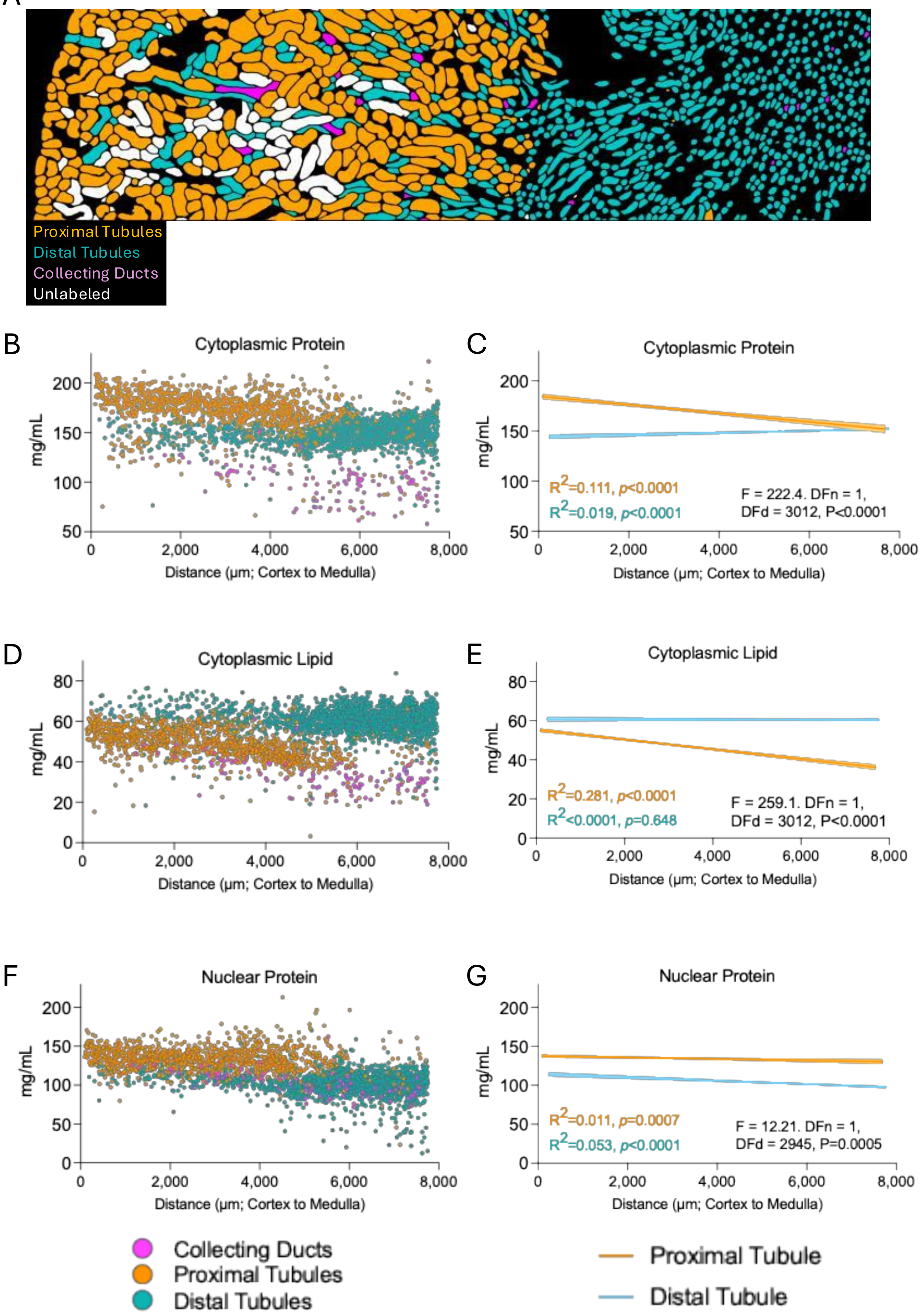

A
Proximal Tubules
Distal Tubules
Collecting Ducts
Unlabeled
B
Cytoplasmic Protein
mg/mL
Distance (μm; Cortex to Medulla)
C
Cytoplasmic Protein
mg/mL
R2=0.111, p<0.0001
R2=0.019, p<0.0001
F = 222.4. DFn = 1,
DFd = 3012, P<0.0001
Distance (μm; Cortex to Medulla)
D
Cytoplasmic Lipid
mg/mL
Distance (μm; Cortex to Medulla)
E
Cytoplasmic Lipid
mg/mL
R2=0.281, p<0.0001
R2<0.0001, p=0.648
F = 259.1. DFn = 1,
DFd = 3012, P<0.0001
Distance (μm; Cortex to Medulla)
F
Nuclear Protein
mg/mL
Distance (μm; Cortex to Medulla)
G
Nuclear Protein
mg/mL
R2=0.011, p=0.0007
R2=0.053, p<0.0001
F = 12.21. DFn = 1,
DFd = 2945, P=0.0005
Distance (μm; Cortex to Medulla)
Collecting Ducts
Proximal Tubules
Distal Tubules
Proximal Tubule
Distal Tubule

**Supp. Fig. 8**

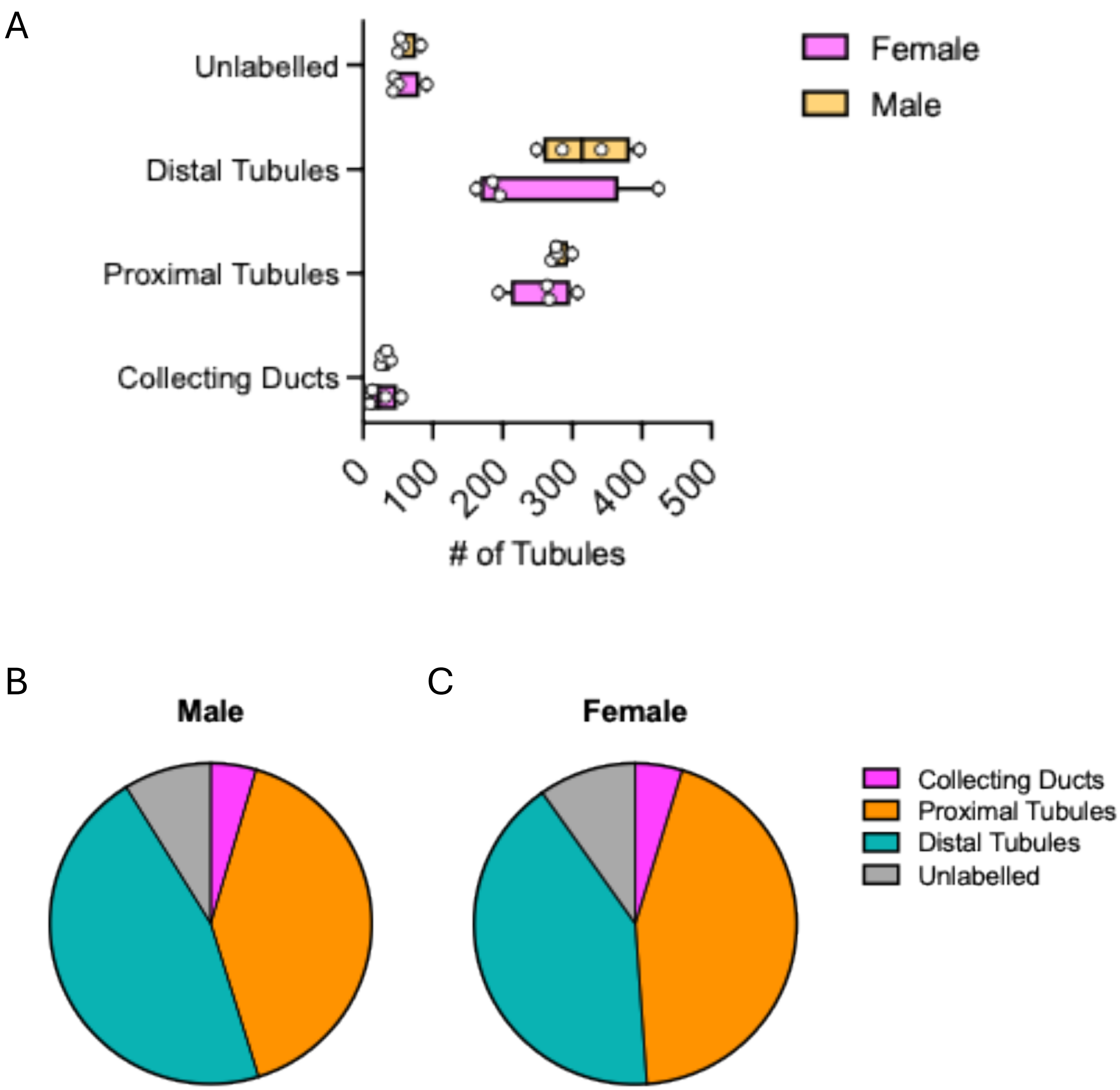
A
Unlabelled
Distal Tubules
Proximal Tubules
Collecting Ducts
0
100
200
300
400
500
### of Tubules
Female
Male
B
Male
C
Female
Collecting Ducts
Proximal Tubules
Distal Tubules
Unlabelled

**Supp. Fig. 9**

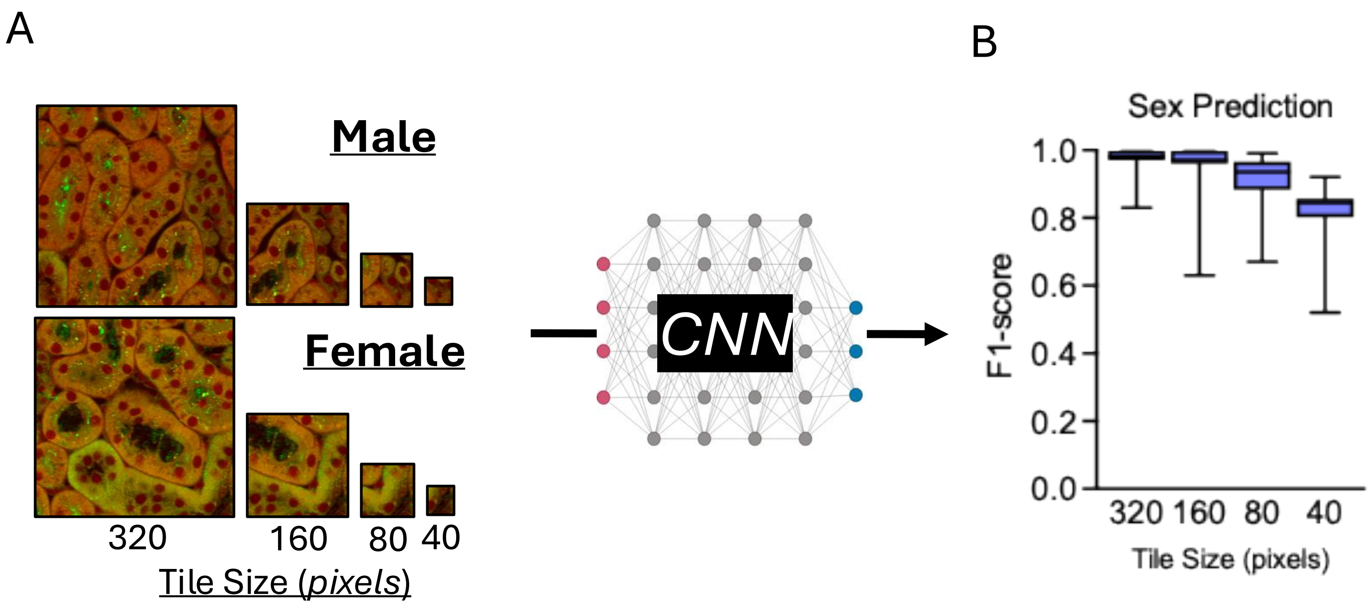

A
Male
Female
320
160
80
40
Tile Size (pixels)
CNN
B
Sex Prediction
1.0
0.8
0.6
0.4
0.2
0.0
F1-score
320 160 80 40
Tile Size (pixels)

Supp. Fig. 10

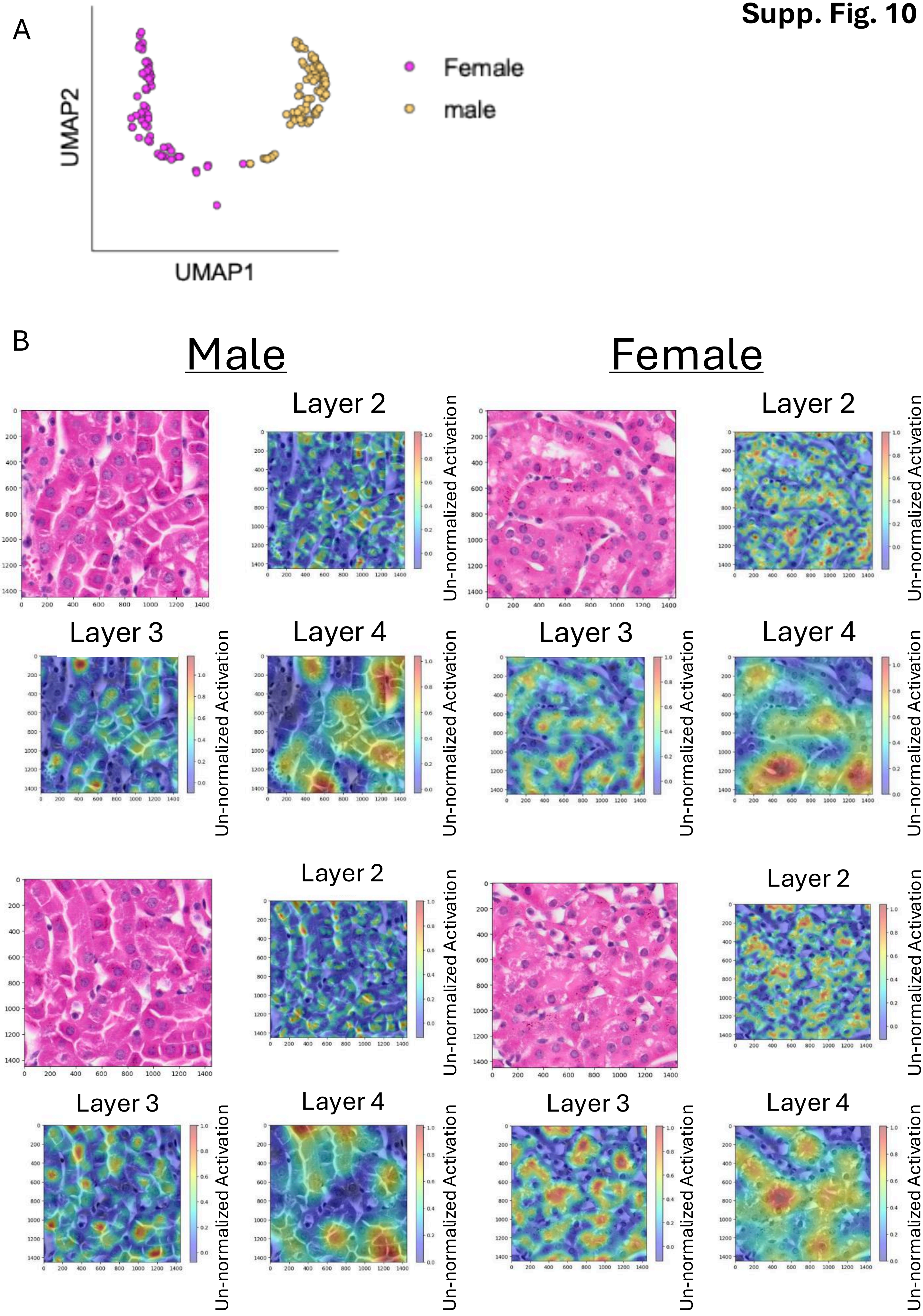

A
UMAP2
UMAP1
Female
male
B
Male
Female
Layer 2
Layer 3
Layer 4
Un-normalized Activation

**Supp. Fig. 11**

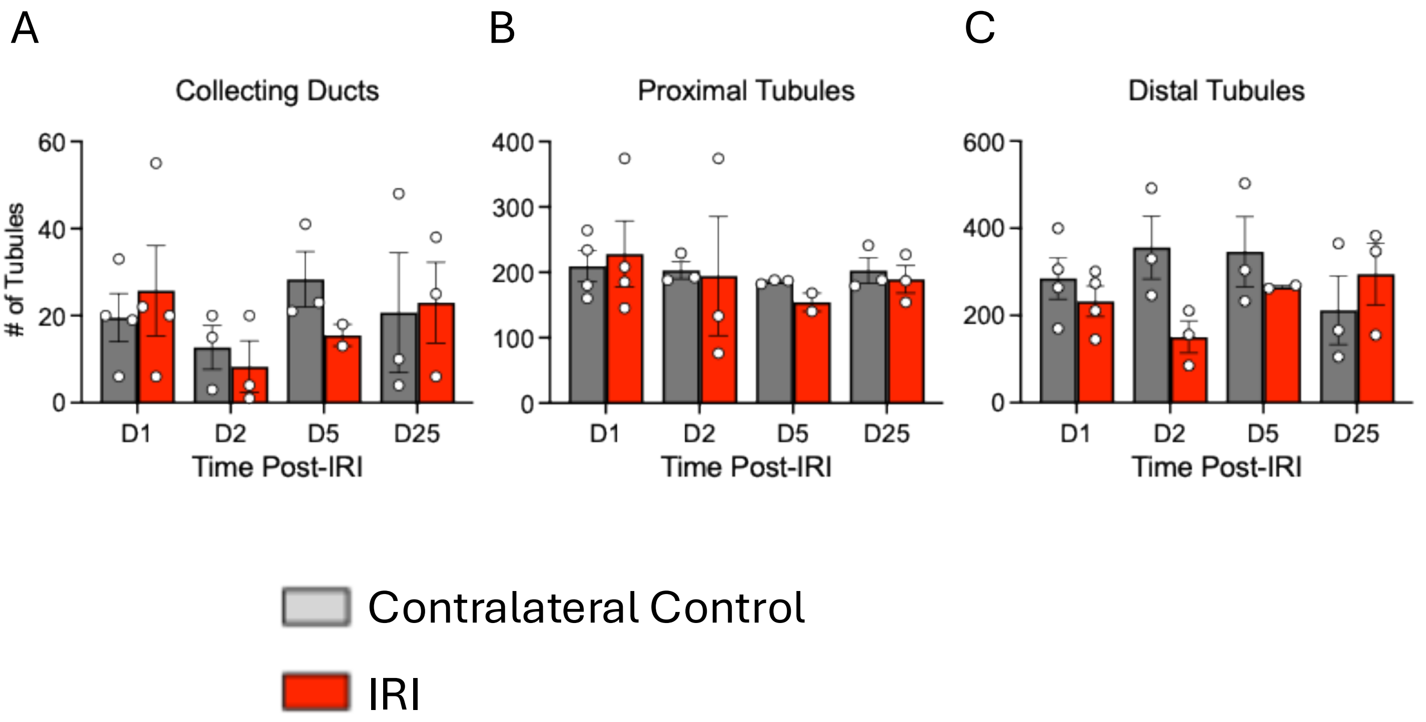

A
Collecting Ducts
### of Tubules
60
40
20
0
D1
D2
D5
D25
Time Post-IRI
B
Proximal Tubules
400
300
200
100
0
D1
D2
D5
D25
Time Post-IRI
C
Distal Tubules
600
400
200
0
D1
D2
D5
D25
Time Post-IRI
Contralateral Control
IRI

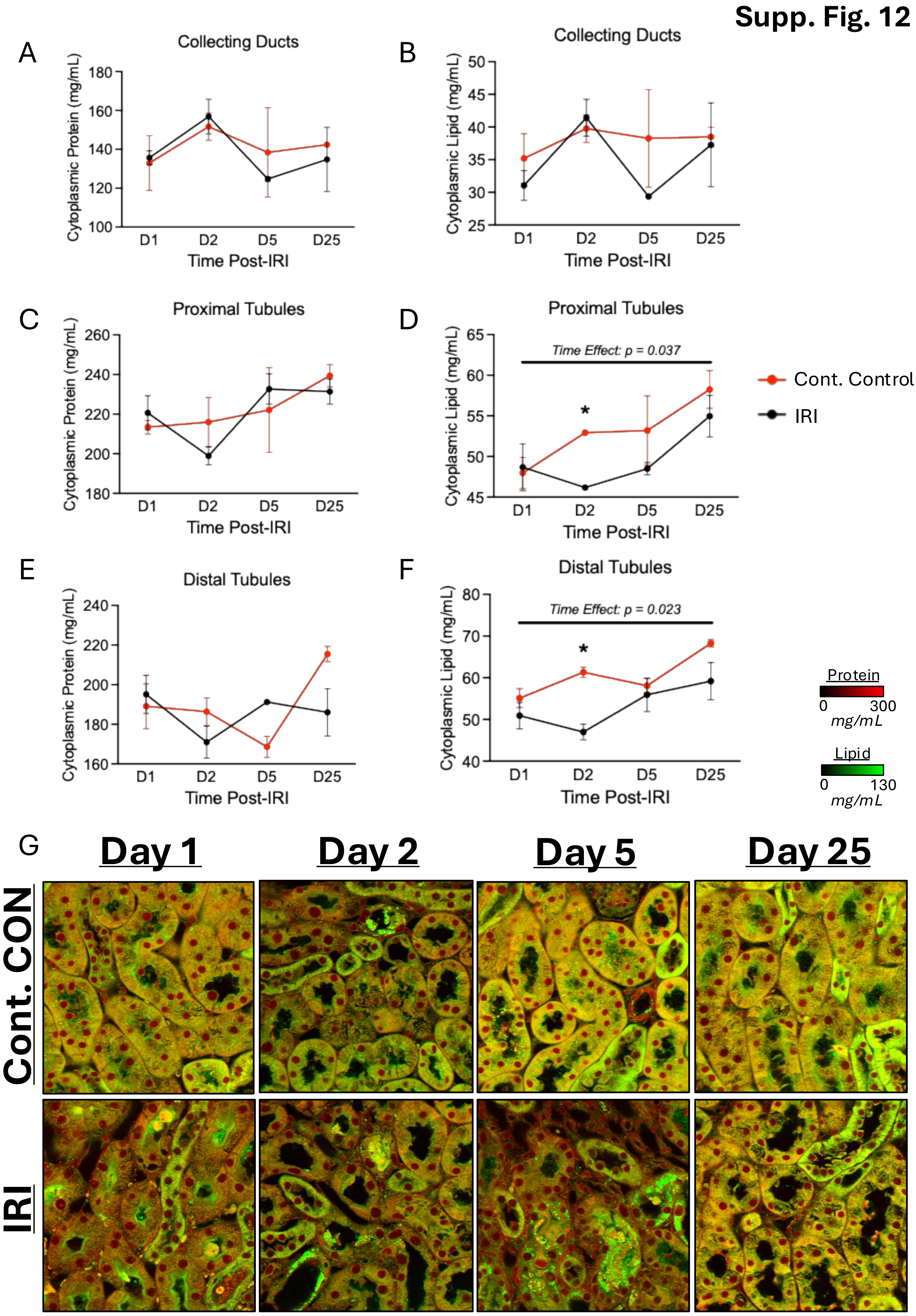

Supp. Fig. 12
A
Collecting Ducts
Cytoplasmic Protein (mg/mL)
180
160
140
120
100
D1
D2
D5
D25
Time Post-IRI
B
Collecting Ducts
Cytoplasmic Lipid (mg/mL)
50
45
40
35
30
25
D1
D2
D5
D25
Time Post-IRI
C
Proximal Tubules
Cytoplasmic Protein (mg/mL)
260
240
220
200
180
D1
D2
D5
D25
Time Post-IRI
D
Proximal Tubules
Time Effect: p = 0.037
*
Cytoplasmic Lipid (mg/mL)
65
60
55
50
45
D1
D2
D5
D25
Time Post-IRI
Cont. Control
IRI
E
Distal Tubules
Cytoplasmic Protein (mg/mL)
240
220
200
180
160
D1
D2
D5
D25
Time Post-IRI
F
Distal Tubules
Time Effect: p = 0.023
*
Cytoplasmic Lipid (mg/mL)
80
70
60
50
40
D1
D2
D5
D25
Time Post-IRI
Protein
0
300
mg/mL
Lipid
0
130
mg/mL
G
Day 1
Day 2
Day 5
Day 25
Cont. CON
IRI

**Supp. Fig. 13**

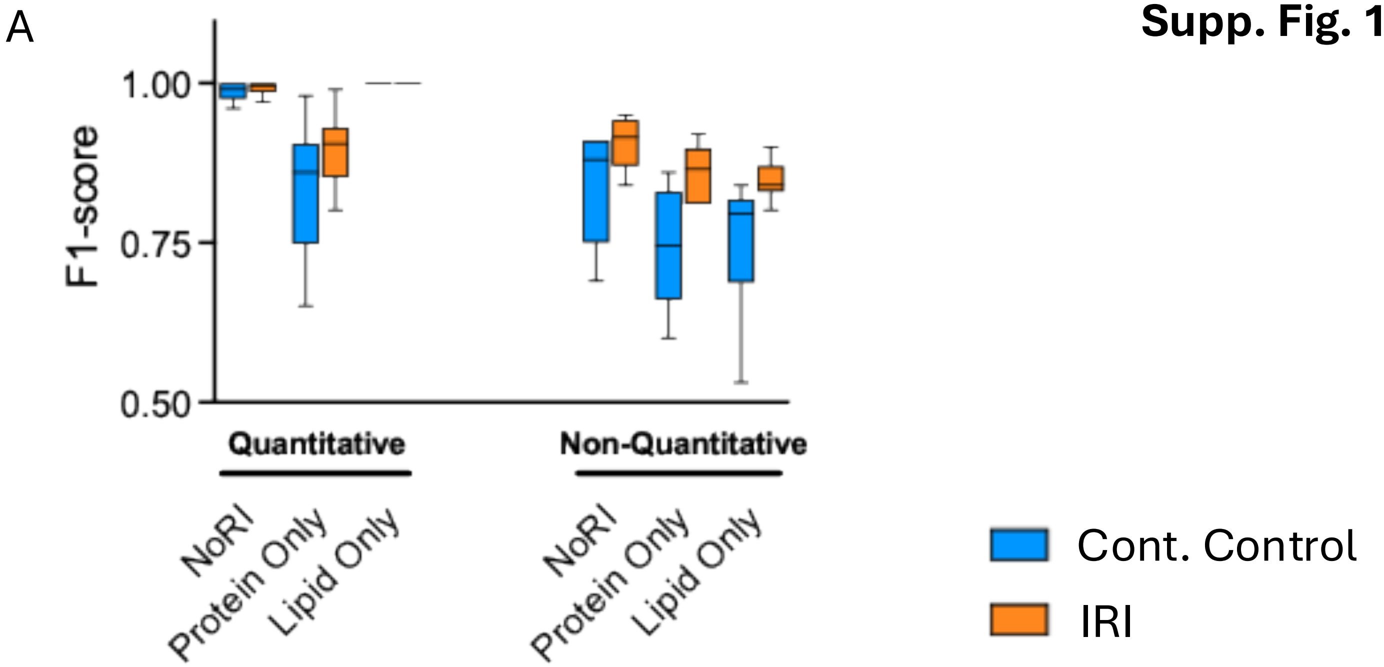
A
F1-score
1.00
0.75
0.50
Quantitative
Non-Quantitative
NoRI
Protein Only
Lipid Only
NoRI
Protein Only
Lipid Only
Cont. Control
IRI

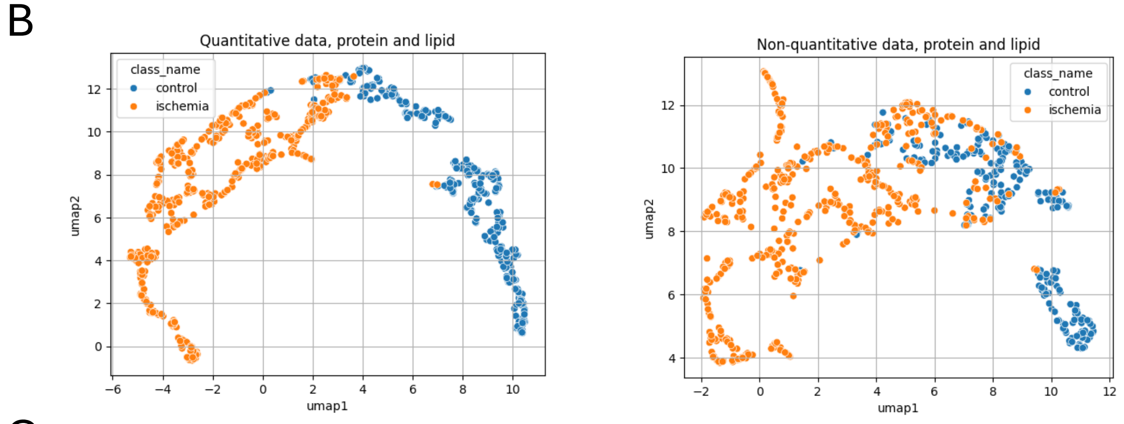
B
Quantitative data, protein and lipid
Non-quantitative data, protein and lipid
class_name
control
ischemia
umap1
umap2

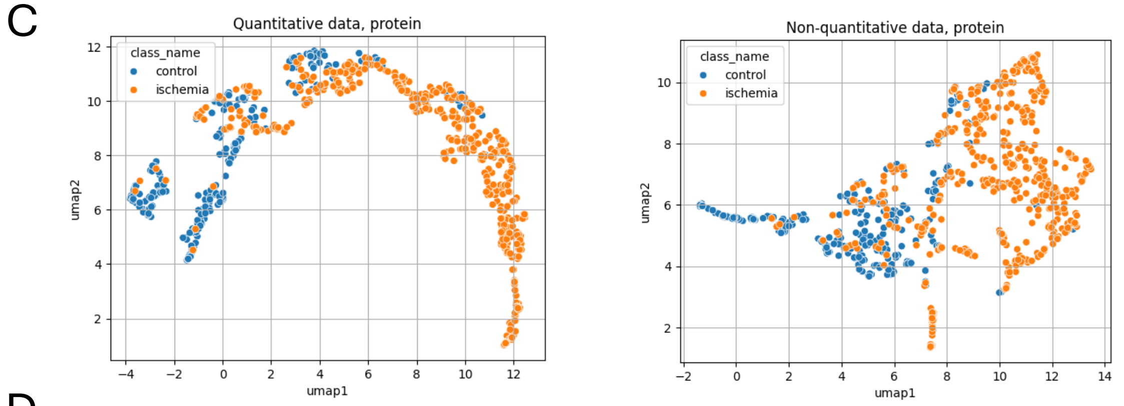
C
Quantitative data, protein
Non-quantitative data, protein
class_name
control
ischemia
umap1
umap2

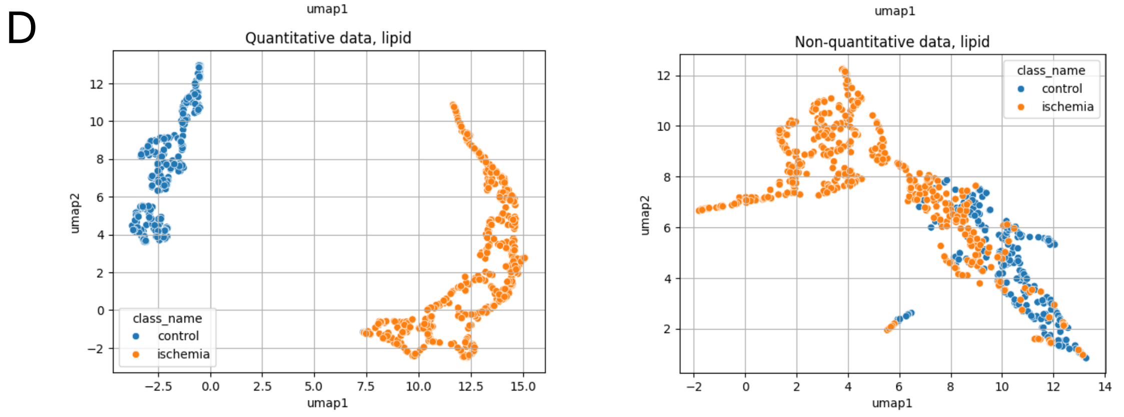
D
Quantitative data, lipid
Non-quantitative data, lipid
class_name
control
ischemia
umap1
umap2

**Supplementary Figure 1.** Representative images and quantitative comparisons by NoRI for different handling conditions required for immunofluorescence purposes prior to imaging samples.

A, Representative NoRI images from all conditions for visual comparison.

B, Scatter plot of tubule cytoplasmic protein and lipid concentration measured by NoRI across each of the experimental handling conditions being tested. Each data point represents one tubule.

C, Violin plots of tubule protein and lipid concentrations across each experimental handling condition.

**Supplementary Figure 2.** Representative image of NoRI slice size and location.

A, Visual representation of the location of NoRI images obtained throughout all experiments. NoRI images are overlaid on top of a brightfield image of the kidney slice they were taken from.

B, Visual representation of four-channel immunofluorescence from a whole kidney slice.

C, LTL+ channel from Panel B.

D, UMOD+ channel from Panel B.

E, ENDO+ channel from Panel B.

F, AQP2+ channel from Panel B.

**Supplementary Figure 3**. Schematic of NoRI image analysis pipeline.

Mouse kidneys of different sexes, or under different experimental conditions, were resected and analysed by two means: convolutional neural network (CNN) analysis (pipeline a) and quantitative analysis of segmented regions of interest (pipeline b). CNN analysis (left and center, pipeline a) was performed by splitting NoRI image tiles (512$\mu m^2$) into 16.5–132.5$\mu m^2$ tiles, with 50% overlap between adjacent tiles. These tiles were used for fine-tuning and validation of models to identify regions of interest and determine biological predictions. Regions that predict biological state with high confidence were subsequently overlaid on segmented masks for labelling and context. From this, specific biological structures identified by CNN analysis were quantified. Basic quantification of anatomic structures (pipeline b) was performed by first segmenting known biological structures in kidney images. Segmentation was performed using a combination of classical image processing, deep learning and machine learning techniques, including the Segment Anything Model (SAM) for tubule segmentation(11). Post-processing steps, such as morphological operations and expert validation, were applied to refine segmentation accuracy. For comparison with SAM, a YOLOv8-based object segmentation model(12) was also fine-tuned to detect kidney tubules in NoRI images using annotated data derived from the protein and lipid signals. From these models, tubules, nuclei, brush border, and nucleoli were parsed out of NoRI images for direct quantification of each structure, including cytoplasmic protein and lipid concentrations (top, right X-Y scatter plot) between tubules, and the nuclear protein and brush border protein and lipid concentrations for all tubules (bottom, right violin plot). These segmented regions are also incorporated into the ResNet50 feature identification pipeline (pipeline a), to assign labels to regions identified by ResNet50 activation maps, as shown here for sex differences in microvasculature protein concentrations (bottom, center violin plot).

**Supplementary Figure 4.** Representative images of nucleoli segmentation from NoRI images. Images are presented as the original NoRI image to the left of each tubule mask (white) within which nuclei (red masks) and nucleoli (green masks) are also segmented. Four different examples are presented.

**Supplementary Figure 5.** Representative view of NoRI, immunofluorescence, and segmentation masks for one kidney cross section.

A, Absolute quantitative greyscale readout of protein content by NoRI. Signal intensity scale bar is presented below the panels. Scale bar in micrometers is presented in the top right corner.

B, Absolute quantitative greyscale readout of lipid content by NoRI. Signal intensity scale bar is presented below. Scale bar in micrometers is presented in the top right corner.

C, Greyscale representation of LTL immunofluorescence. Signal intensity scale bar is presented below. Scale bar in micrometers is presented in the top right corner.

D, Greyscale representation of UMOD immunofluorescence. Signal intensity scale bar is presented below. Scale bar in micrometers is presented in the top right corner.

E, Greyscale representation of AQP2 immunofluorescence. Signal intensity scale bar is presented below. Scale bar in micrometers is presented in the top right corner.

F, Greyscale representation of Endomucin immunofluorescence. Signal intensity scale bar is presented below. Scale bar in micrometers is presented in the top right corner.

G, Tubule mask produced by SAM on NoRI images presented in Panels A and B. For display purposes each tubule is eroded by a circular structuring element with radius 5. Scale bar in micrometers is presented in the top right corner.

H, Nuclei mask produced by YOLO on NoRI images presented in Panels A and B. Scale bar in micrometers is presented in the top right corner.

I, Brush border mask produced by YOLO on NoRI images presented in Panels A and B. Scale bar in micrometers is presented in the top right corner.

J, Lumen mask produced by YOLO on NoRI images presented in Panels A and B. Scale bar in micrometers is presented in the top right corner.

**Supplementary Figure 6.** Analysis of protein and lipid concentrations by anatomical region.

A, Scatter plot of tubule cytoplasmic protein and lipid concentration measured by NoRI split by anatomical region. Each datapoint represents an individual tubule, color coded by the anatomical region they were derived from.

B, Violin plots of cytoplasmic protein and lipid concentrations measured by NoRI, split by tubule type as defined by immunofluorescence positivity of one of three proteins, between anatomical regions. Dashed lines represent group median, dotted lines represent $25^{th}/75^{th}$ confidence intervals. Coll. Duct = Collecting Ducts; Prox. = Proximal Tubules; Dist. = Distal Tubules.

C, Cohen's *d* effect size's between anatomical regions, split by tubule type, from data presented in Panel B. Coll. Duct = Collecting Ducts; Prox. = Proximal Tubules; Dist. = Distal Tubules; Rost. = Rostral pole; Caud. = Caudal pole; Mid. = Midpole.

**Supplementary Figure 7.** Anatomical analysis of nuclear protein and cytoplasmic protein and lipid concentrations from Cortex to Medulla.

A, Representative tubule mask image cross-section from cortex to medulla. Tubules are color coded according to immunofluorescence identification of tubule type.

B, Scatter plot of cytoplasmic protein concentration measured by NoRI as a product of spatial location from cortex to medulla. Each datapoint represents an individual tubule, color coded by tubule identity.

C, Summary of linear regression analysis from data in Panel B for proximal and distal tubule cytoplasmic protein from cortex to medulla. Data were analysed by simple linear regression. Trend lines are shown with 95% confidence bands of the line of best fit displayed. Regression coefficients are displayed for each tubule type, and the summary of differences in slopes and intercepts are also displayed.

D, Scatter plot of cytoplasmic lipid concentration measured by NoRI as a product of spatial location from cortex to medulla. Each datapoint represents an individual tubule, color coded by tubule identity.

E, Summary of linear regression analysis from data in Panel D for proximal and distal tubule cytoplasmic lipid from cortex to medulla. Data were analysed by simple linear regression. Trend lines are shown with 95% confidence bands of the line of best fit displayed. Regression coefficients are displayed for each tubule type, and the summary of differences in slopes and intercepts are also displayed.

F, Scatter plot of nuclear protein concentration measured by NoRI as a product of spatial location from cortex to medulla. Each datapoint represents an individual tubule, color coded by tubule identity.

G, Summary of linear regression analysis from data in Panel D for proximal and distal tubule nuclear protein from cortex to medulla. Data were analysed by simple linear regression. Trend lines are shown with 95% confidence bands of the line of best fit displayed. Regression coefficients are displayed for each tubule type, and the summary of differences in slopes and intercepts are also displayed.

**Supplementary Figure 8.** Cumulative tubule counts and proportions in male and female kidneys.

A, Cumulative tubule counts, split by tubule type, for male and female mice. Each datapoint represents the number of tubules for an individual mouse. Boxes represent group median, with $25^{th}/75^{th}$ percentiles. Whiskers represent min-to-max values.

B, Proportional representation of each tubule type across all male kidney samples.

C, Proportional representation of each tubule type across all female kidney samples.

**Supplementary Figure 9.** Tile-size comparison for ResNet50 prediction accuracy by F1-score in male and female mouse kidneys.

A, Visual representation of tile sizes input into ResNet50 for tubule type identification in male and female mouse kidneys.

B, Box-whisker plot of F1-scores for tubule type prediction accuracy across different input tile sizes. Boxes represent median with $25^{th}/75^{th}$ percentiles. Whiskers represent min-to-max values.

**Supplementary Figure 10.** Analysis of H&E stained kidney sections from male and female kidneys by ResNet50.

A, UMAP plot of male and female tubules analysed in H&E stained images by ResNet50.

B, Male and Female visual representations of image tiles analysed by ResNet50. Each raw H&E image tile is presented with multi-color activation heatmaps representing layers 2–4 of the ResNet50 analysis pipeline—that is each iteration of the filter set for feature identification (where layers 2 to 4 represents fine to gross regions within a given image tile)—that distinguish male and female kidneys. Scale bars represent un-normalised activation scores.

**Supplementary Figure 11.** Cumulative tubule counts at each time point post-IRI for Contralateral control and IRI, split by tubule type.

A, Cumulative counts of Collecting ducts at each time point post-IRI. Each datapoint represents the number of tubules at each time point for a given mouse. Bars represent group mean. Error bars represent Standard Error of the Mean.

B, Cumulative counts of proximal tubules at each time point post-IRI. Each datapoint represents the number of tubules at each time point for a given mouse. Bars represent group mean. Error bars represent Standard Error of the Mean.

C, Cumulative counts of distal tubules at each time point post-IRI. Each datapoint represents the number of tubules at each time point for a given mouse. Bars represent group mean. Error bars represent Standard Error of the Mean.

**Supplementary Figure 12.** Quantification of tubule protein and lipid concentration by NoRI at each time point post-IRI for Contralateral control (Cont. Control) and IRI.

A, Line graph of collecting duct cytoplasmic protein measured by NoRI at each time point post-IRI. Each point represents the group mean for $n$=2–4 mice per time point, per group. Error bars represent Standard Error of the Mean.

B, Line graph of collecting duct cytoplasmic lipid measured by NoRI at each time point post-IRI. Each point represents the group mean for $n$=2–4 mice per time point, per group. Error bars represent Standard Error of the Mean.

C, Line graph of proximal tubule cytoplasmic protein measured by NoRI at each time point post-IRI. Each point represents the group mean for $n$=2–4 mice per time point, per group. Error bars represent Standard Error of the Mean.

D, Line graph of proximal tubule cytoplasmic lipid measured by NoRI at each time point post-IRI. Each point represents the group mean for *n*=2–4 mice per time point, per group. Error bars represent Standard Error of the Mean.

E, Line graph of distal tubule cytoplasmic protein measured by NoRI at each time point post-IRI. Each point represents the group mean for *n*=2–4 mice per time point, per group. Error bars represent Standard Error of the Mean.

F, Line graph of distal tubule cytoplasmic lipid measured by NoRI at each time point post-IRI. Each point represents the group mean for *n*=2–4 mice per time point, per group. Error bars represent Standard Error of the Mean.

G, Representative NoRI images for CON and IRI at each time point post-IRI.

**Supplementary Figure 13**. Case study analysis of the contribution of the quantitative power of NoRI protein and lipid data.

A, F1 scores produced by ResNet50 on NoRI images comparing the ability to distinguish control from ischemic kidneys on day 2 post-reperfusion injury between protein and lipid NoRI data versus protein or lipid data in isolation. Data for each condition is split into fully quantitative (all samples normalized together prior to analysis by ResNet50) and non-quantitative (each image normalized in isolation, and each protein and lipid channel also normalized in isolation prior to ResNet50 analysis).

B, UMAP plot of control and ischemic kidney tubules on day 2 post-reperfusion injury from quantitative (left) and non-quantitative (right) NoRI images containing both protein and lipid data analysed by ResNet50.

C, UMAP plot of control and ischemic kidney tubules on day 2 post-reperfusion injury from quantitative (left) and non-quantitative (right) NoRI images containing protein data only analysed by ResNet50.

D, UMAP plot of control and ischemic kidney tubules on day 2 post-reperfusion injury from quantitative (left) and non-quantitative (right) NoRI images containing lipid data only analysed by ResNet50.
