## Supplementary Table 1 for "Normalized Raman Imaging for Studies of Tissue Physiology of the Kidney"

**Supplementary Table 1.** Geometric features used by ResNet50 with descriptive overviews.

| Feature | Description |
|---|---|
| Area | The total number of pixels enclosed within the contour. |
| Perimeter | The length of the contour boundary. |
| Aspect Ratio | The ratio of the contour's width to its height, as determined by its bounding box. |
| Extent | The ratio of the contour area to the area of its bounding box. |
| Solidity | The ratio of the contour area to the area of its convex hull, indicating how filled the contour is. |
| EquivDiameter | The diameter of a circle that has the same area as the contour. |
| Orientation | The angle at which the contour is oriented, typically calculated from an ellipse fitted around the contour. |
| Major Axis Length | The length of the longest axis of an ellipse fitted around the contour. |
| Minor Axis Length | The length of the shortest axis of an ellipse fitted around the contour. |
| Eccentricity | A measure of how elongated the contour is, calculated as the ratio of the distance between the ellipse's foci to the length of the major axis. |
| Convex Hull | The smallest convex shape that can fully enclose the contour. |
| Compactness | The ratio of the perimeter squared to the area, providing an indication of how compact the shape is. |
| Rectangularity | The ratio of the contour area to the area of its minimum enclosing rectangle. |
| Roundness (Circularity) | A measure of how closely the contour resembles a perfect circle. |
| Elongation | A measure of how stretched the contour is, often calculated as the ratio of the major axis length to the minor axis length. |
| Nuclei Number | The number of predicted nuclei inside the contour. |
| Minimal Distance and Average Distance | The average and minimal distance from the nuclei within the contour to the contour boundary, often used to assess nuclear distribution. |
| Minimal Distance and Average Distance between nuclei | The average and minimal distance between nuclei. |
